# Weber’s law of proportional processing influences coevolution of ornaments and preferences in models of sexual selection

**DOI:** 10.64898/2026.05.01.722204

**Authors:** Kathryn Bullough, Laura A. Kelley, Bram Kuijper

## Abstract

Mate preferences are often influenced by the magnitude of sexual signals, which are presumed to indicate underlying aspects of signaller quality. Although the perception of these signals depends on sensory processes, the role of perceptual adaptations and constraints in mate assessment is frequently overlooked. Many sensory systems follow Weber’s law of proportional processing, where discrimination between signals is based upon their proportional, or relative, difference rather than their absolute difference. Because preference strength varies with relative trait magnitude, Weber’s law could strongly influence sexual selection, changing the coevolution of traits and preferences. Here, we explore the consequences of Weber’s law for sexual selection using individual-based models, applying Scalar Utility Theory to mate choice. We investigate the coevolution of male ornaments and female preferences under both Fisherian and good genes scenarios, as well as scrutinizing the sexual selection of multiple ornaments and preferences. Including Weber’s law in these models either reduced ornament exaggeration, or promoted exaggeration and diversification of ornaments and preferences, depending on the costs of choice and how rapidly female survival decreases when preferences evolve away from the naturally selected optimum. These results highlight the importance of perception and cognitive processing in shaping sexual selection and its evolutionary impacts.

## Introduction

Sexual selection is responsible for some of the most elaborate ornaments and behaviours across the natural world, causing rapid evolution across taxa, ecosystems, and sensory modalities, and even leading to speciation (Rosenthal, 2017; Stafstrom & Hebets, 2013). Numerous attempts have been made to model this process and explore how selection that generally only acts on half the population can generate such rapid evolution and drive ornament values well beyond naturally selected optima (Kuijper et al., 2012; Rosenthal, 2017). Following the most general model of Fisherian sexual selection using choosers (generally females) and signallers (generally males) by Lande (1981), a variety of extensions have been developed incorporating aspects such as direct or indirect benefits for choosers (Hamilton & Zuk, 1982; Hoelzer, 1989; Iwasa & Pomiankowski, 1999; Kirkpatrick & Barton, 1997; Møller & Jennions, 2001; Pomiankowski et al., 1991), biased mutations (Pomiankowski et al., 1991), costly female preferences (Iwasa et al., 1991), and signalling efficacy (Tazzyman et al., 2014).

Although these models employ many different approaches and include a vast array of necessary assumptions, there is one previously unrecognised assumption that they all have in common – a unit increase in an ornament has the same effect across the whole ornament space. When it comes to the sensory processes involved in mate choice, namely how female sensory systems process information acquired from male signals, we find that this assumption is rarely, if ever, true. One well-documented mechanism that animals (including humans) use to compare stimuli is through proportional processing, or Weber’s law (Akre & Johnsen, 2014; Bullough et al., 2023; Fechner, 1966; Weber & Mollon, 1978). Weber’s law states that stimuli are compared based on their proportional (relative) difference rather than absolute difference in magnitude. Consequently, Weber’s law implies that as the magnitude of a stimulus increases, it becomes more difficult to detect a given difference in magnitude (Akre & Johnsen, 2014; Dehaene, 2003). It is described mathematically as:

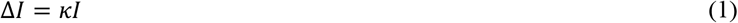

Δ*I* is the minimum detectable change in a stimulus of magnitude *I*, and *κ* is the constant required for a just noticeable difference, often termed the Weber fraction. Weber’s law underlies how signals are processed during mate choice in a broad range of taxa and sensory modalities, for example when females process the male ‘chuck’ calls in Túngara frogs (*Engystomops pustulosus*) (Akre et al., 2011) or the male body area in green swordtail fish (*Xiphophorus hellerii*) (Caves & Kelley, 2023).

A key consequence of Weber’s law during mate choice is that, as male signal magnitudes increase, it may become more difficult for receivers to differentiate between signallers (Cohen, 1984). Weber’s law could therefore have implications for the evolution of sexual ornaments and preferences for them. For example, as ornament size increases, female ability to detect absolute differences between them becomes increasingly difficult, and ornaments would evolve to ever-larger magnitudes for the difference to be detectable to females, potentially resulting in exaggerated chase-away selection (Akre & Johnsen, 2014). Alternatively, sexual selection for larger ornaments may weaken with increasing ornament size, due to female inability to discriminate between high magnitude ornaments (Akre & Johnsen, 2014; LaBarbera et al., 2020; Nachev et al., 2017). One potential outcome of weaker selection on ornament size is that it could favour redirecting investment into other signal elements. Hence, a central prediction is that Weber’s law is more likely to result in multidimensional signals (e.g., Rowe, 1999; van Doorn & Weissing, 2004) relative to conventional models of mate choice (Akre & Johnsen, 2014; p.66 in Rosenthal, 2017). To our knowledge, however, there have been no attempts to formally assess these predictions by reconciling Weber’s law with influential models on the evolution of mate choice.

Here we take the first steps to investigate how including a female Weber-based preference function in models of sexual selection could influence predicted outcomes for the coevolution of male ornaments and female preferences. We consider the evolution of single and multiple ornaments and preferences in both Fisherian and good genes contexts.

To implement an evolutionary model of mate choice in the context of Weber’s law, we draw on Scalar Utility Theory (SUT), which uses Weber’s law to describe how stimuli of different magnitudes are represented in memory (Gibbon, 1977; Gibbon et al., 1988; Kacelnik & Brito e Abreu, 1998; Rosenström et al., 2016). At the heart of SUT is that the representation of a stimulus of magnitude *t* in an animal’s memory has a distribution that is normal with a mean *t* and a standard deviation that is proportional to the mean: *σ* × *t*. The original inception of SUT focused on memory of time perception (Gibbon, 1977). However, since then SUT has been used to assess decision making in a broader range of contexts (Bateson & Kacelnik, 1995a, 1995b; Brito e Abreu & Kacelnik, 1999; Kacelnik & Bateson, 1996; Reboreda & Kacelnik, 1991, and others), successfully predicting that perception of the difference between two stimuli is characterized by noise proportional to the absolute magnitude of the stimuli (e.g., Figure 5 in Bateson & Kacelnik, 1995a; Kacelnik & Brito e Abreu, 1998).

Applying SUT to the context of mate choice implies that females decide on which mate to accept by retrieving a sample value for the different observed males from memory, so that ornaments of a larger magnitude are perceived from memory with larger levels of noise. Doing so mirrors central ideas in the context of Weber’s law, that when stimulus magnitudes are higher, more discrimination errors will be made (Worsley et al., 2025). However, we are currently lacking formal predictions on how such magnitude-dependent discrimination errors affect the evolution of costly female preferences for male ornaments. Here, we do so by allowing for magnitude-dependent discrimination errors to influence mating decisions when females evaluate single vs multiple ornaments through an open-ended preference function, which are conventionally used in the majority of sexual selection models.

### Model 1: Fisherian sexual selection

Following Fawcett et al. (2007) and Fawcett et al. (2011), an evolutionary individual-based model of sexual selection was developed. Following birth, individuals go through a round of survival selection (natural selection), mate choice, and sexual reproduction (in which offspring inherit mutated versions of their parents’ traits), before the next generation fully replaces the previous one and the cycle starts again. This life cycle was repeated for 150,000 generations. Models were developed for both unidimensional sexual selection (where females express a preference for only one male ornament, and only a single preference-ornament pair is present) and multidimensional sexual selection (where females express preferences for multiple male ornaments, and multiple preference-ornament pairs are present).

#### Survival selection

The individuals in our model represent a sexually reproducing, diploid population of a total number of *N* = 5000 of males and females at a 1:1 sex ratio. Following classical models of sexual selection (Mead & Arnold, 2004), we consider the evolution of two sets of diploid gene loci: a vector **T** = (*T*_*i*_) of male ornament loci and a vector **P** = (*P*_*i*_) of female preference loci. The subscript *i* reflects that our model allows for *n* evolving pairs of genetically unlinked ornament and preference loci, allowing us to compare the effect of Weber’s law in unidimensional (*n* = 1) versus multidimensional (*n* > 1) contexts (e.g., Iwasa & Pomiankowski, 1994; Pomiankowski & Iwasa, 1993; van Doorn & Weissing, 2004). In line with these quantitative genetic models of sexual selection, we assume that ornament and preference loci show continuous variation, where ornament and preference alleles can be any real number. Following the seminal model by Pomiankowski and Iwasa (1993), a value of *P*_*i*_ < 0 indicates preference for smaller values of ornament *T*_*i*_, whilst *P*_*i*_ > 0 indicates preference for bigger values of ornament *T*_*i*_. Females with *P*_*i*_ = 0 mate at random with respect to the value of ornament *T*_*i*_. Whilst both sexes carry the genes for ornament and preference loci (no sex-linked inheritance is assumed), only males express the ornament and only females express the preference (Mead & Arnold, 2004; but see Albert & Otto, 2005). Following previous models and for the sake of tractability, we ignore environmental variation in the expression of *P*_*i*_ and *T*_*i*_, although the modelling framework can be extended to include this.

The survival optimum for the male ornament phenotype was set at a value of 0 (i.e. using a hypothetical fish example, representing a tail length that optimises survival performance), with the male survival probability Pr(male survives) decreasing on either side of this optimum, according to the function:

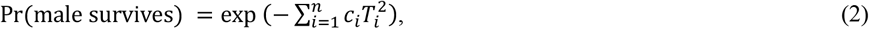

where *c*_*i*_ is a non-negative constant scaling the cost of the ornament (following Pomiankowski & Iwasa, 1993, equation 10a). Female preferences are also likely to be costly (Pomiankowski, 1987), so we also assume a decrease in female survival away from the naturally selected optimum *P*_*i*_ = 0, according to the function (following Pomiankowski & Iwasa, 1993, sequation 10b):

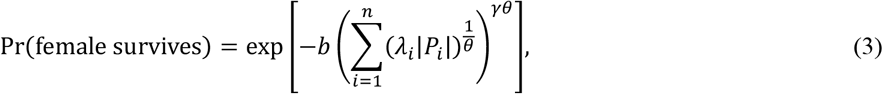

where *b* is a nonnegative constant scaling the cost of choice, and *λ*_*i*_ describes the relative costliness of the *i*th female preference relative to the other female preference loci. Next, the parameter *θ* describes the joint cost of choice: when *θ* tends towards 0, expressing multiple preferences does not greatly increase the cost of choice compared to a unidimensional model of sexual selection. By contrast, as *θ* tends towards 1, costs of choice combine in a supermultiplicative manner so that expressing multiple preferences considerably increases the cost of choice over and above that in a unidimensional model of sexual selection. Finally, *γ* is the power cost coefficient of female preference (determining the shape of the decline of female fitness around the optimum due to *b*).

#### Mate choice

After natural selection, each surviving female then chooses a mate. To do so, each female randomly samples 10 individuals among the surviving males (each male can mate multiple times). The female chooses one of the 10 males based on the odds *ψ*(**T**|**P**) that a sampled male with ornament vector **T** will be chosen by female with preference vector **P**:

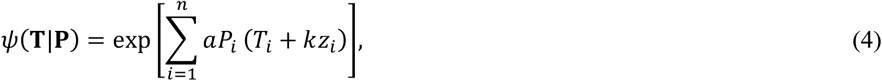

where *a* is a nonnegative constant scaling the importance of the male ornament to female choice (e.g., Lande, 1981); *k* is a nonnegative constant that reflects how strongly Weber preferences influence mate choice, equivalent to the Weber fraction in eq. (1) above. When *k* = 0, Weber preferences are absent, so that males will be chosen based on open-ended preferences, which is the preference function of choice in nearly all models of sexual selection (following Lande, 1981, equation 8a). However, a corollary of open-ended preferences is that a difference in ornament value of a fixed magnitude Δ always results in same mating advantage, regardless of the overall magnitude of *T*_*i*_ (see supplement S1 for a proof). Hence, this precludes any notion of Weber’s law, so that the case of *k*=0 serves as a null model in which Weber preferences are absent.

For scenarios in which the constant *k* > 0, however, the preference function in eq. (4) now captures the essential element of the Weber process, namely that the observational noise increases with the size of the ornament, making it more difficult to accurately perceive the relative size of this ornament the larger it becomes (Gibbon, 1977; Kacelnik & Brito e Abreu, 1998; Rosenström et al., 2016). We do so by sampling a random number *z*_*i*_ from a normal distribution with mean 0 and a standard deviation *σT*_*i*_ that is a scalar function of the overall size of the ornament *T*_*i*_ and where *σ* is the standard deviation of a standard normal distribution with value *σ* = 1.

#### Reproduction and inheritance

Following mate choice, *N* newborn offspring were randomly sampled from the mating pairs. Each pair produces a clutch size of 10 offspring, inheriting alleles in a Mendelian fashion. For each trait, mutations occur with probability *μ*_*P,i*_ and *μ*_*T,i*_ per allele per generation for loci *P*_*i*_ and *T*_*i*_ respectively. For the preference loci, the value of the *P*_*i*_ allele then changes by an increment *z*_*P*_ drawn from a uniform probability distribution U(−0.4,0.4). For the ornament loci, we follow previous models of sexual selection by including biased mutations, so that mutations are more likely to decrease the value of the ornament rather than to increase it. Previous models have shown that biased mutations result in stable exaggeration of ornaments and preferences, even when female preferences are costly (Andersson, 1994; Pomiankowski et al., 1991). To include biased mutations on the ornament, let *u*_*t*_ reflect the probability that a mutational increment *z*_*T*_ sampled from a uniform distribution U(0,0.4) changes the value of a *T*_*i*_ allele by a negative amount (−*z*_*T*_), while 1 − *u*_*t*_ is the probability that the value of *T*_*i*_ allele is increased by a positive amount (*zT*). We consider scenarios where *u*_*t*_ ≥ 0.5.

After reproduction, all adult individuals are replaced by *N* individuals sampled from the offspring generation with offspring sex being randomly determined (generations are nonoverlapping) after which the cycle continues.

### Model 2: Good-genes sexual selection

In the good-genes model, an extra genetic locus is included: a single diploid heritable quality locus *v*, carried and expressed by both sexes and located on a different chromosome than *T* and *P* loci. Similar to previous models (Iwasa & Pomiankowski, 1994; Iwasa et al., 1991), this locus influences survival to maturity. Both *T*_*i*_ and *v* determine the phenotype of the male ornament in the good genes model, according to the function

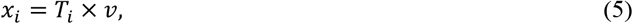

where *x*_*i*_ denotes the *i*th ornament’s realised phenotype. This reflects a condition-dependent indicator (Iwasa et al., 1991), because males in better condition can grow more exaggerated ornaments than males in poorer condition with the same genetic value for the ornament *T*_*i*_.

#### Survival selection

Crucially, in line with existing models of condition-dependent indicators (Andersson, 1994; Iwasa et al., 1991), the realised size of the ornament *x*_*i*_ affects both survival and mating success. Our model can, however, be easily extended to consider other forms of condition-dependence, such as epistatic and revealing indicators (Iwasa et al., 1991). In the case of condition-dependent indicators, a male’s survival probability is then given by:

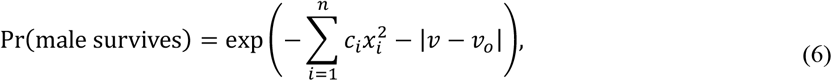

where *v*_*o*_ is the optimum value of *v* for survival (following Iwasa & Pomiankowski, 1994, equations 2a & 13). Female survival decreases on either side of the naturally selected optimum for female preferences (*P*_*i*_ = 0) according to the function:

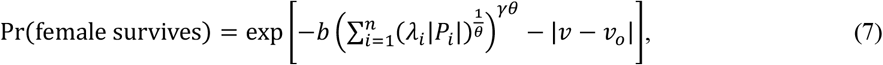

(following Iwasa & Pomiankowski, 1994, equations 3 & 13)). See the analogous eq. (3) on Fisherian sexual selection above for definitions of the other parameters.

#### Mate choice

Similar to the model on Fisherian sexual selection above, each surviving female then chooses a mate out of a random sample of 10 surviving males. This time, mate choice is based on the condition-dependent indicator *x*_*i*_, so that the odds *ψ*(***x***|**P**) that a sampled male with ornament vector ***x*** will be chosen by female with preference vector **P** are given by (see also eq. [4]):

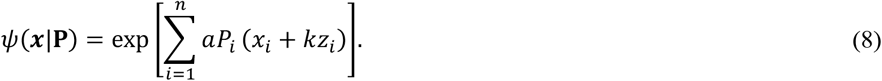

Again, this allows us to have a case of *k* = 0 in which this preference function serves as a null model where Weber preferences are absent. Scenarios in which the constant *k* > 0 captures the essential element of the Weber process, namely that the observational noise increases with the size of the ornament *x*_*i*_, making it more difficult to accurately perceive the size of this ornament the larger it becomes (see eq. [4]). Again, *z*_*i*_ reflects a number drawn from a normal distribution with mean 0 and a standard deviation *σx*_*i*_ that is a scalar function of the overall size of the ornament *x*_*i*_ and where *σ* is the standard deviation of a standard normal distribution with value *σ* = 1.

#### Reproduction and inheritance

As in the Fisherian sexual selection model, following mate choice we randomly sample *N* newborn offspring from the pairs of surviving females and mates.

Mutation on the quality allele occurs with probability *u*_*v*_ (*u*_*v*_ = 0.05 throughout this study), causing the allelic value to change by an increment *z*_*v*_ drawn from a uniform probability distribution U[0,0.8]. The increment is negative (−*z*_*v*_) with probability *u*_*v*_ and positive otherwise, reflecting a mutation bias as in standard models of good-genes sexual selection. Following previous models of good-genes sexual selection (e.g., van Doorn and Weissing (2006); Iwasa and Pomiankowski (1994); van Doorn and Weissing (2004)) male ornaments *T*_*i*_ are not subject to negative mutation bias in the good-genes model (*u*_*t*_ = 0.5).

Each mating produces a clutch size of 10 offspring, inheriting alleles at all loci *T*_*i*_, *P*_*i*_, and *v* in a Mendelian fashion. After reproduction all adult individuals die and are replaced by *N* individuals sampled from offspring generation, with offspring sex being randomly determined (generations are nonoverlapping) after which the cycle continues.

Simulations were implemented in C++, and the code is publicly available at [https://github.com/kathrynbullough/Weber].

## Results

### Fisherian sexual selection

#### Unidimensional models

Male ornaments and female preferences coevolve in similar ways as predicted by analytical models of sexual selection when Weber preferences are absent (*k* = 0). For example, similar to classical models of sexual selection (Pomiankowski et al., 1991), stable exaggeration of ornaments and preferences increases in trait value in the presence of biased deleterious mutations on the ornament (Figure 1A). By contrast, exaggeration away from the naturally selected optima of ornaments and preferences is much more limited in the absence of biased deleterious mutations (*u*_*t*_ = 0.5). When we consider scenarios involving Weber preferences (*k* > 0), exaggeration of ornaments and preferences is notably lower in trait value (Figure 1B, C, D), with an increase in deleterious mutation bias causing a less pronounced increase in the sexually-selected values of *t* and *p*. For resulting variances and covariances between ornaments and preferences, see Figure S3.1, and for varying strengths of the cost of choice parameter *b*, see Figures S3.2 & S3.3).

**Figure 1:**
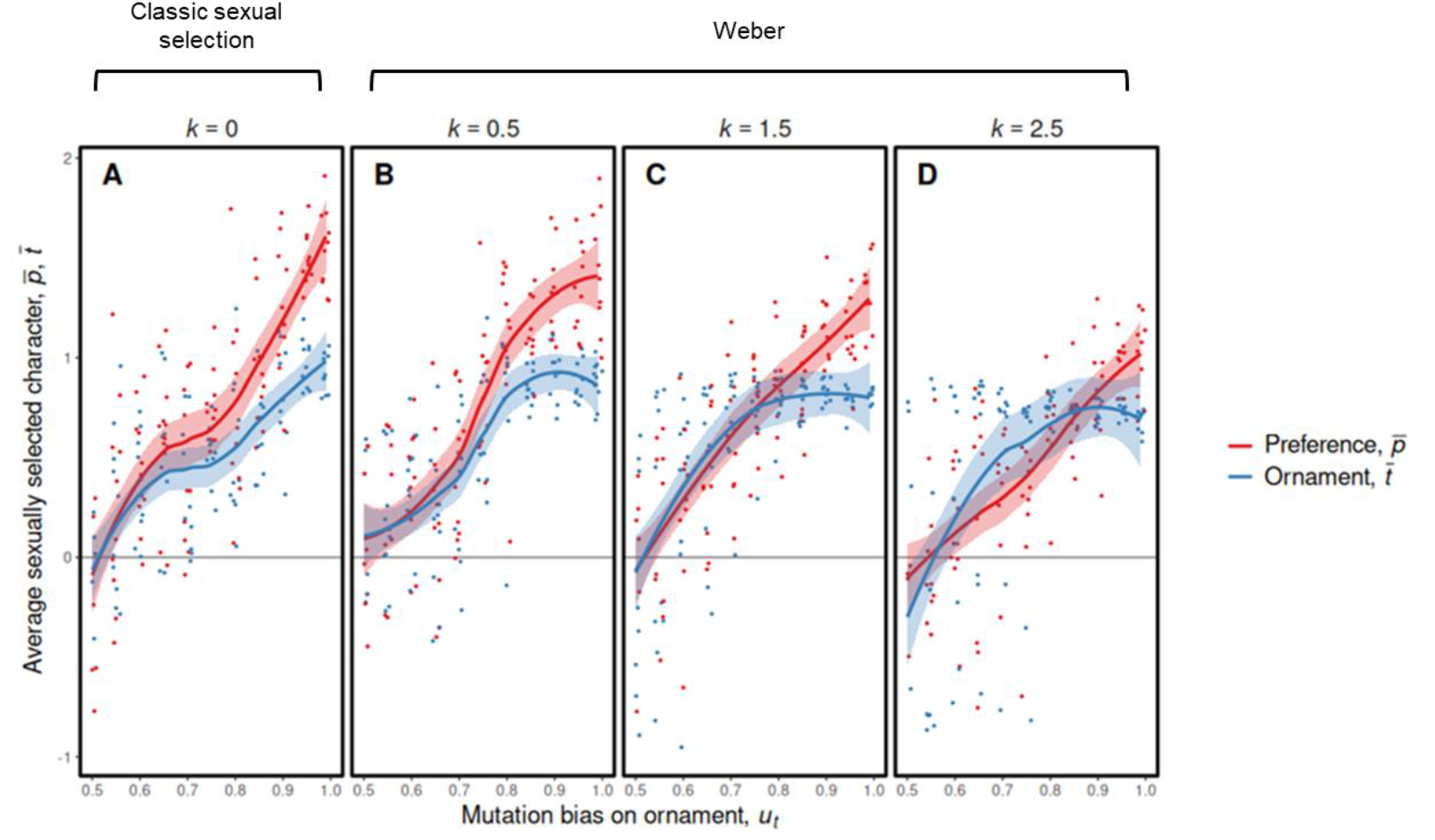
Varying the strength of Weber preferences in a univariate model of Fisherian sexual selection with costs to female preference. As the deleterious mutation bias *u*_*t*_ on the male ornament increases, exaggeration of ornaments and preferences also increases, yet with increasing noise *k* reflecting Weber preferences exaggeration of ornaments and preferences moderately reduces. Parameters: *a*=1, *λ*=1, *b*=0.0025, *c*=0.5, *μ*_*P*,_ and *μ*_*T*,_ =0.05, *γ*=2, and 10 replicate simulations for each unique parameter combination. Dots represent the endpoints of each individual simulation, and the starting values for ornamentation and preference were *t*=1 and *p*=3. Variances and covariances are depicted in Figure S3.1.

#### Multidimensional models

We then focused on a scenario in which we coevolved two ornaments and two preferences (Figure 2). Each ornament is affected by a mutation bias of *u*_*t*_ = 0.99, which we expect to result in the stable exaggeration of each ornament and preference pair. Indeed, similar to unidimensional models of Fisherian sexual selection, Figures 2A-C show that multiple ornaments and preferences in the presence of open-ended preferences (*k* = 0) result in stable exaggeration away from the naturally selected optimum. Moreover, for larger values of γ (reflecting how strongly survival costs of each preference increase when away from its naturally selected optimum), we find that exaggeration of ornaments and preferences is reduced, similar to classical models of multiple sexual preferences (Pomiankowski & Iwasa, 1993).

**Figure 2:**
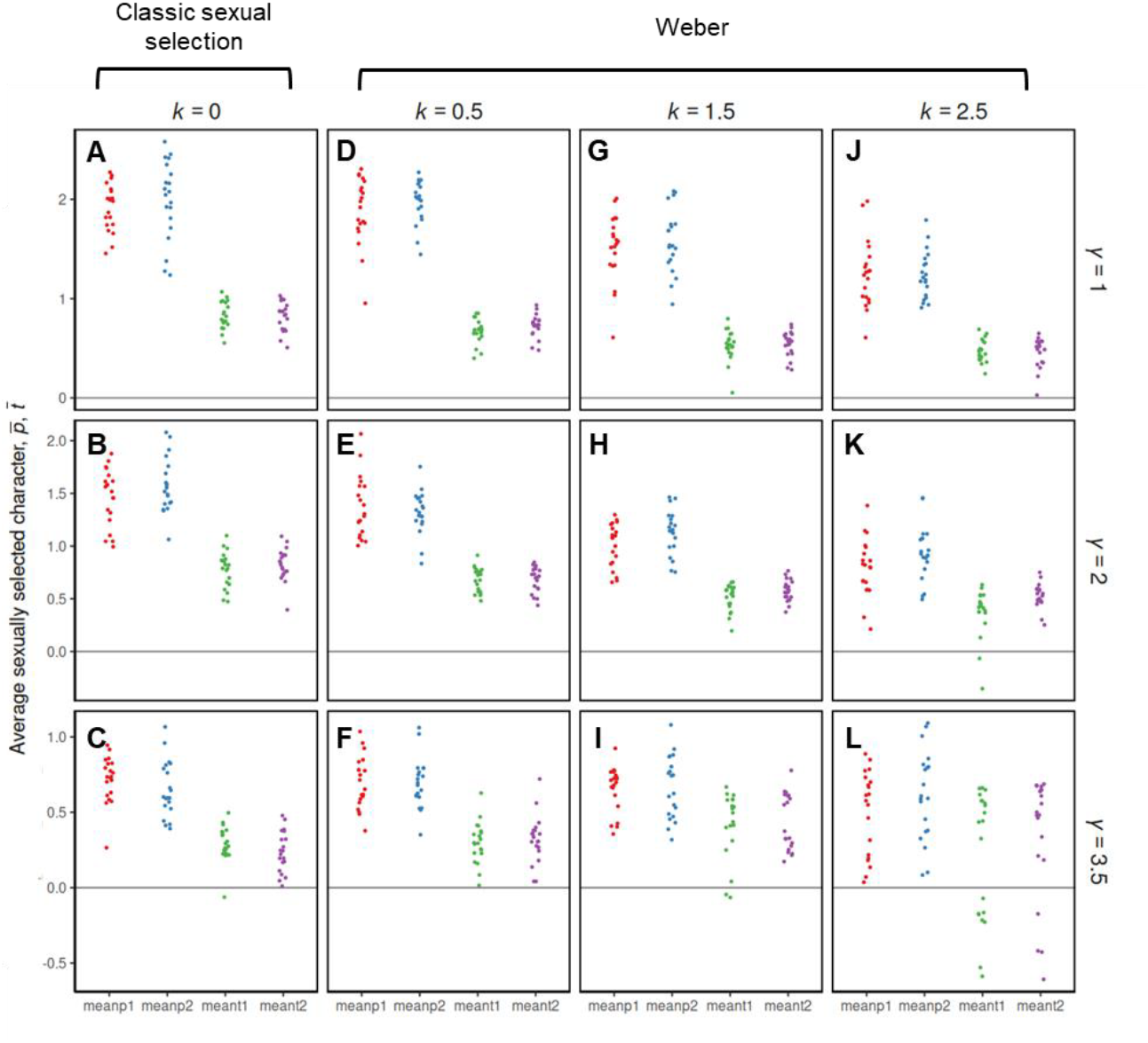
A multidimensional individual-based model of Fisherian sexual selection, showing equilibrium exaggeration of two male ornaments (*t*_1_ and *t*_2_) and two female preferences (*p*_1_ and *p*_2_) at generation 150,000. Exaggeration is less pronounced under Weber preferences for low values of γ, whilst under high values of γ ornaments and preferences appear to diverge with Weber preferences. For the models shown, *a*=1, *λ*=1, *b*=0.0025, *c*=0.5, *μ*_*P*,_ and *μ*_*T*,_ =0.05, *ϑ*=0.2, *u*_*t*_ = 0.99, and each set of parameters was replicated 20 times. Dots represent the endpoints of each individual simulation, and the starting values for ornamentation and preference were *t*_*1*_=*t*_*2*_=1 and *p*_*1*_=*p*_*2*_=3. Variances and covariances are depicted in Figure S3.4).

When we consider scenarios involving Weber preferences (*k* > 0), exaggeration of ornaments and preferences still occurs, but is notably reduced relative to classical open-ended preference models (Figure 2D-L, ‘Weber’ versus Figure 2A-C ‘open-ended’). For example, for *k* = 2.5, the average values of both ornaments are ∼40% reduced in magnitude relative to *k* = 0. This large reduction in ornament exaggeration due to the presence of Weber preferences generalizes across increasing values of γ (Figure 2J-L vs Figure 2A-C), or when the expression of multiple preferences reduces survival in a submultiplicative versus supermultiplicative ways (as measured by the parameter θ, see Figure S3.5).

To assess the prediction that Weber preferences may result in diversification towards a larger number of sexually selected traits, we varied the number *n* of evolving ornament-preference pairs well beyond *n* = 2 (Figure 3). Unsurprisingly, when increasing *n*, ornament exaggeration is reduced, since the costs of expressing ever more ornaments substantially reduces a male’s survival. Figures 3C-H show that the relationship between *n* and ornament exaggeration is qualitatively similar for Weber vs open-ended preferences, but overall exaggeration is more reduced for Weber preferences relative to open-ended preferences. Consequently, our model predicts that Weber preferences result in reduced exaggeration of multiple sexually selected traits, relative to open-ended preferences. Again, for larger values of γ, we find that exaggeration of ornaments and preferences is reduced.

**Figure 3:**
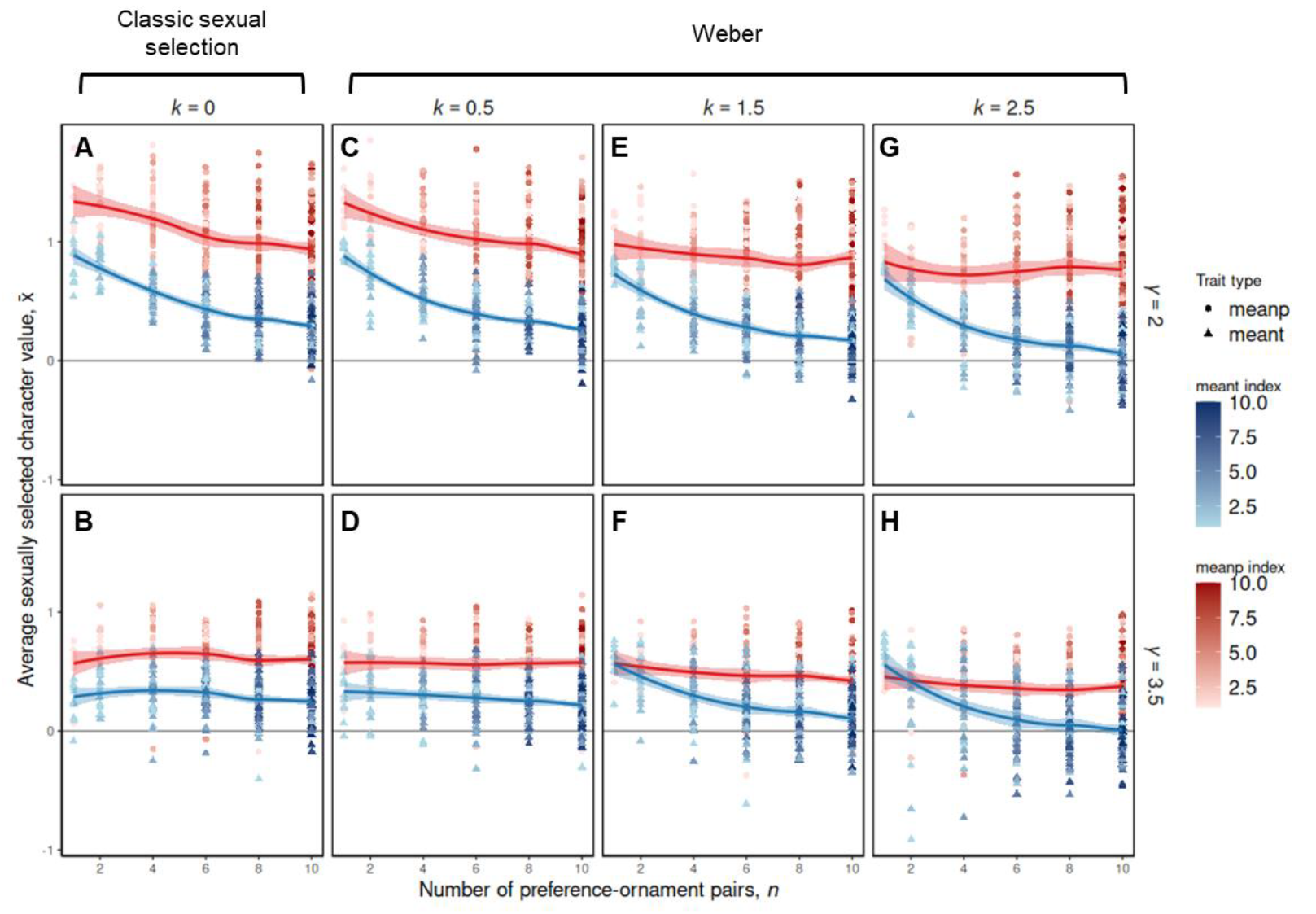
Varying the number of evolving preference-ornament pairs (*n*) and its effect on exaggeration of individual ornaments and preferences in the context of Fisherian sexual selection, under varying values of γ (reflecting how strongly survival costs of each preference increase when away from its naturally selected optimum). Panels A&B: classic open-ended preferences, panels C-H: Weber preferences for *k* = 0.5, *k* = 1.5 and *k* = 2.5. Unsurprisingly, overall exaggeration of individual ornaments and preferences is reduced when increasing *n*, due to the increased survival costs that are incurred by expressing increasing numbers of ornaments and preferences. Interestingly, however, average exaggeration of preferences stabilises for increasing *n*. For the models shown, *a*=1, *λ*=1, *b*=0.0025, *c*=0.5, *μ*_*P*,_ and *μ*_*T*,_ =0.05, *ϑ*=0.2, *u*_*t*_ = 0.9, and each set of parameters was replicated 10 times. Dots represent the endpoints of each individual simulation, and the starting values for ornamentation and preference were *t*=1 and *p*=3. Each data point depicts the grand mean value of *p* (and *t*), which reflects the average value taken over all mean values 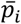 (or 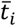) at generation 150,000.

### Good-genes sexual selection

#### Unidimensional models

Under open-ended preferences (*k* = 0), male ornaments and female preferences coevolve as predicted by standard analytical models of good genes sexual selection (e.g., Iwasa et al., 1991), in which exaggerated ornaments and preferences are more likely when the probability of biased mutations reducing male quality (*v*) increases (Figure 4A). Note that exaggerated preferences and ornaments that signal genetic quality appear to settle on two alternative equilibria (either positive or negative, see Figure 4) whereas in Fisherian sexual selection a single, positive equilibrium is typically found (Figures 1-3). This difference stems from assumptions made by classical models regarding mutation biases acting on genetic quality (good genes) versus ornament alleles (Fisher). We further discuss this difference between good genes and Fisher in supplementary section S2, but despite these positive and negative equilibria, qualitative outcomes are similar, with exaggeration being more likely for larger values of the bias *u*_*v*_.

**Figure 4:**
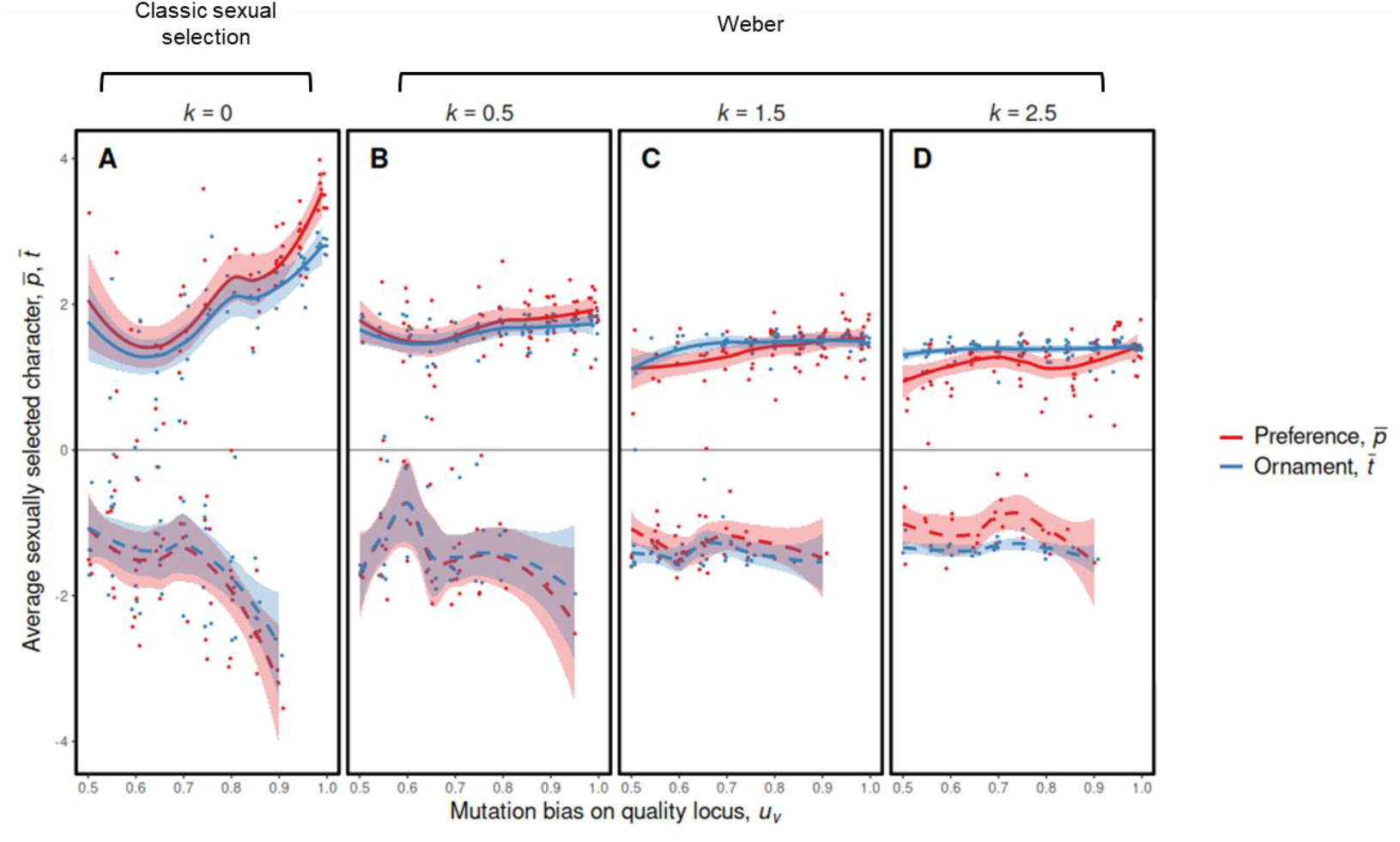
Unidimensional models of good-genes sexual selection, showing the exaggeration of one male ornament (*t*) and one female preference (*p*) for open-ended preferences (panel A) and for different amounts of noise reflecting Weber preferences (panels B, C, D). For all scenarios, as deleterious mutation bias on the male quality locus increases, exaggeration of ornaments and preferences also increases, however this is less pronounced under Weber preferences. For the models shown, *a*=1, *λ*=1, *b*=0.0025, *c*=0.5, *μ*_*P*,_ and *μ*_*T*,_ =0.05, *γ*=2, and each set of parameters was replicated 10 times. Dots represent the endpoints of each individual simulation, and the starting values for ornamentation and preference were *t*=1 and *p*=3. Trendlines and 95% confidence intervals are displayed for both positive and negative exaggeration of characters. Corresponding variances and covariances can be found in Figure S3.6.

When we consider models that involve Weber preferences (*k* > 0), exaggeration of ornaments and preferences is slightly lower relative to *k* = 0, similar to our findings in Fisherian sexual selection (shown in Figure 1). However, we notice that for progressively larger values of *k* the effect of mutation biases becomes less apparent, with stable exaggeration occurring even when biases are absent (Figures 4B, C, D). For resulting variances and covariances between ornaments and preferences, see Figure S3.6, and for varying strengths of the cost of choice parameter *b*, see Figures S3.7 & S3.8).

#### Multidimensional models

Similar to multidimensional models in the context of Fisherian sexual selection (see Figure 2), we assess the exaggeration of ornaments and preferences in a multivariate good-genes model by considering a probability of biased mutation of *u*_*v*_ = 0.99, in which stable exaggeration is expected to occur. We assess the effect of Weber preferences for different costs of maintaining multiple preferences, as measured by the parameters γ (reflecting the curvature with which natural selection increases away from the naturally selected optimum) and θ, which reflects whether natural selection expressing multiple increases in submultiplicative versus supermultiplicative ways (for θ, see Figure S3.10).

We first focus on a scenario of exaggeration in the case of open-ended preferences (*k* = 0). Figure 5B reflects a case similar to the unidimensional model in Figure 4, in which survival selection against female preferences falls away from the naturally selected optimum *p* = 0 in a quadratic fashion (γ = 2): we again find stable exaggeration for both ornaments and preferences away from the naturally selected optimum. Similar to Fisherian sexual selection in Figure 2, we see that exaggeration is reduced for increasing values of γ (e.g., Figure 5A, C) and similar patterns occur for the parameter θ, Figure S3.10A-C.

**Figure 5:**
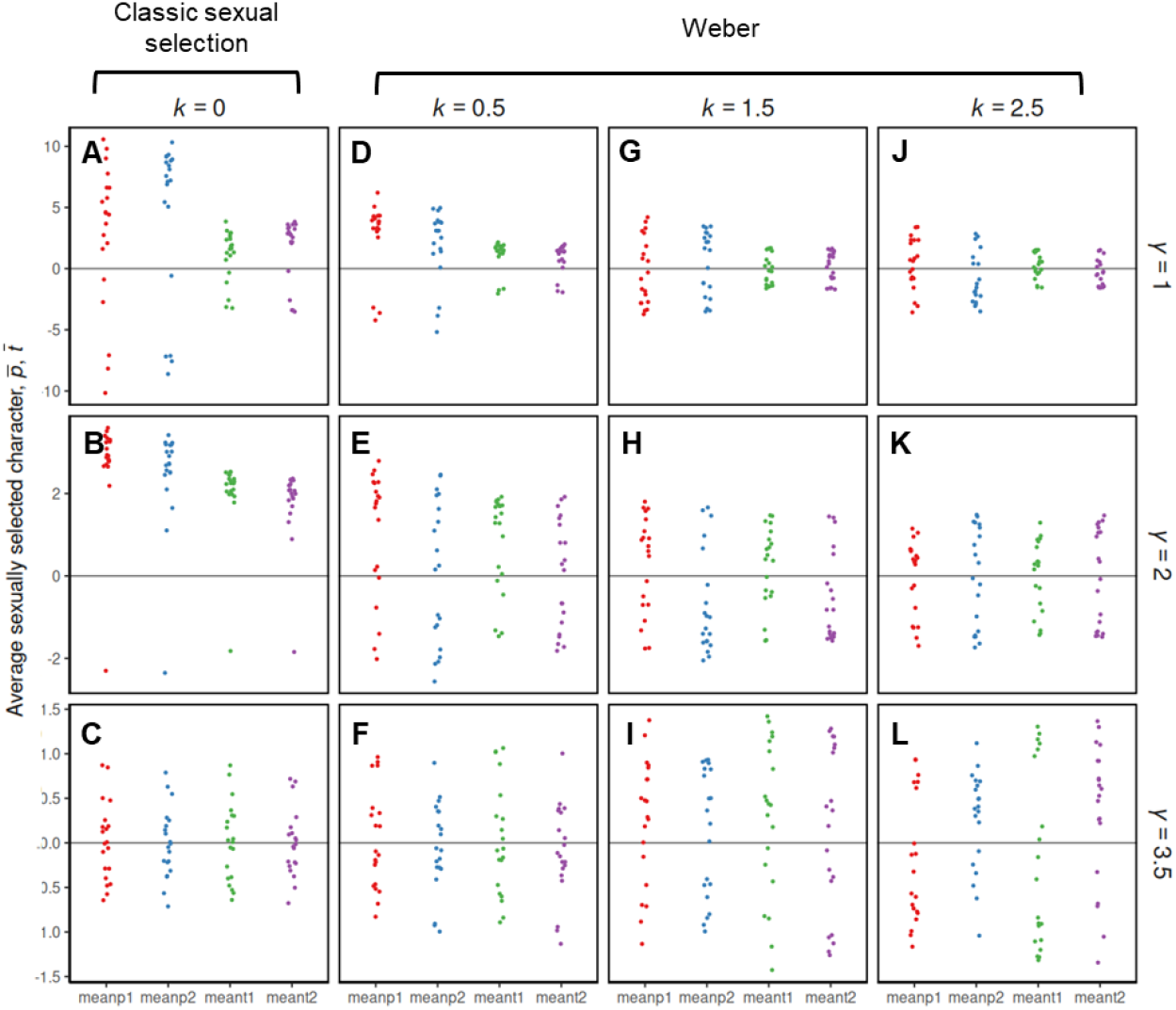
A multidimensional model of good genes sexual selection, showing exaggeration of two male ornaments (*t*_*1*_ and *t*_*2*_) and two female preferences (*p*_*1*_ and *p*_*2*_) under varying values of γ (reflecting how strongly survival costs of each preference increase when away from its naturally selected optimum) at generation 50,000, for a scenario in which biased mutations reducing genetic quality occur with probability *u*_*v*_ = 0.99. Exaggeration is less pronounced under Weber preferences for low values of *γ*, whilst under high values of *γ* ornaments and preferences appear to exaggerate and diverge with Weber preferences. For the models shown, *a*=1, *λ*=1, *b*=0.0025, *c*=0.5, *μ*_*P*,_ and *μ*_*T*,_ =0.05, *ϑ*=0.2, and each set of parameters was replicated 20 times. Dots represent the endpoints of each individual simulation, and the starting values for ornamentation and preference were *t*_*1*_=*t*_*2*_=1 and *p*_*1*_=*p*_*2*_=3. For corresponding variances and covariances, see Figure S3.9.

When we consider the models that involve Weber preferences (*k* > 0) (Figure 5D-L, ‘Weber’), we find again that this generally results in reduced exaggeration relative to scenarios in which *k* = 0 (e.g., compare Figures 5J vs 5A, or 5K vs 5B). An exception occurs, however, for larger values of *γ* (e.g., *γ* = 3.5) in which Weber preferences allow for larger exaggeration in comparison to *k* = 0 (compare Figure 5L vs 5C). Consequently, while lower exaggeration is expected under Weber preferences in most cases, for highly nonlinear costs of female preferences (as measured by *γ*), other outcomes are possible.

To assess the prediction that Weber preferences may result in diversification towards a larger number of sexually selected traits, Figure 6 varies the number *n* of evolving ornament-preference pairs well beyond *n* = 2. Similarly to the Fisher models, increasing *n* reduces ornament exaggeration since expressing ever more ornaments substantially reduces a male’s survival. Again, when looking at smaller values of γ, overall exaggeration is more reduced for Weber preferences relative to open-ended preferences, predicting that Weber preferences result in reduced exaggeration of multiple sexually selected traits (Figures 6A vs 6C, E&G). For larger values of *γ*, Weber preferences allow for larger exaggeration in comparison to *k* = 0, until *n* becomes large, in which case the levels of exaggeration are similar and stabilize around the same values (Figures 6B vs 6D, F&H).

**Figure 6:**
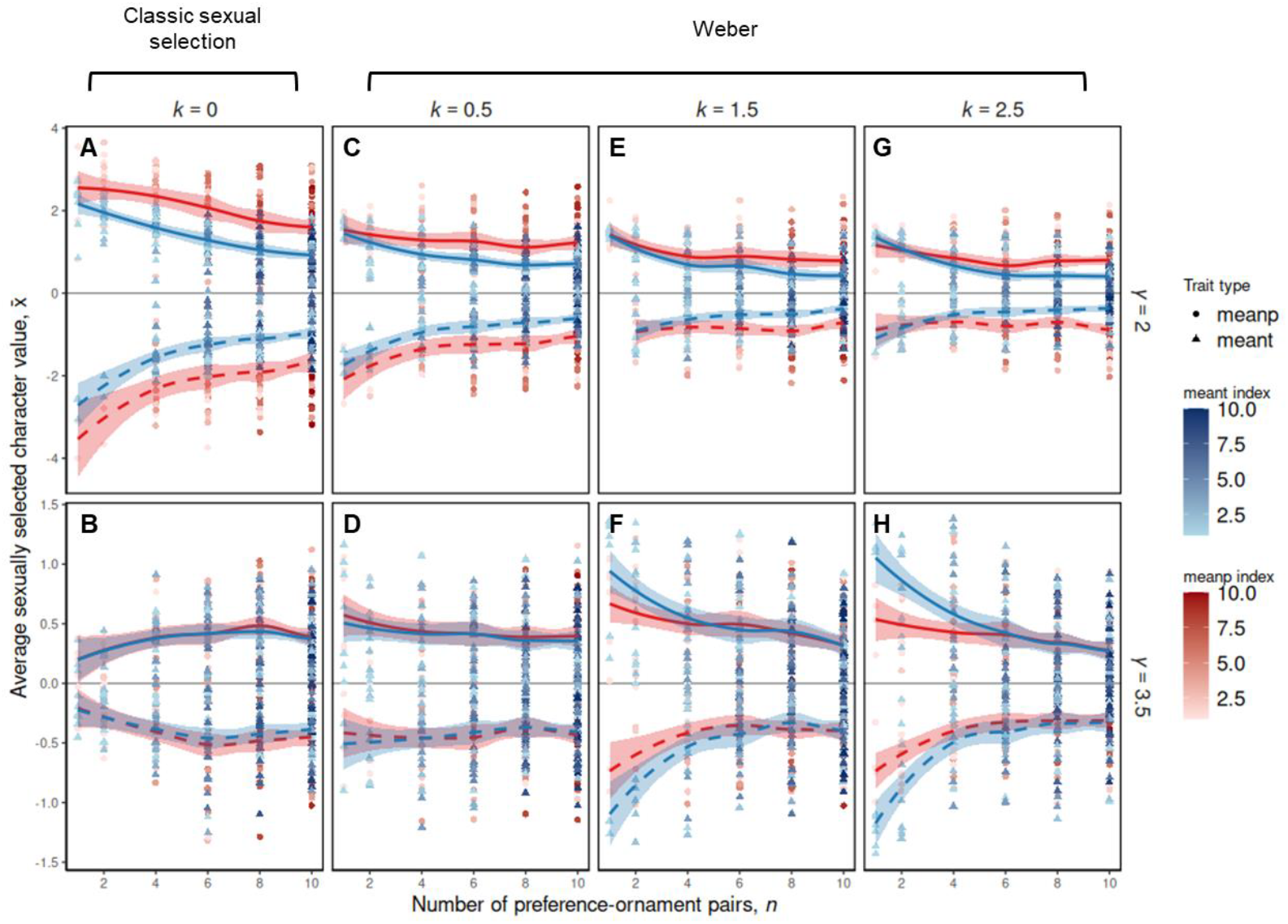
Increasing the number of evolving preference-ornament pairs (*n*) and its effect on exaggeration of individual ornaments and preferences in the context of good genes sexual selection, under varying values of γ (reflecting how strongly survival costs of each preference increase when away from its naturally selected optimum). Panels A&B: classical open-ended preferences, panels C-H: Weber preferences for *k* = 0.5, *k* = 1.5 and *k* = 2.5. Unsurprisingly, overall exaggeration of individual ornaments and preferences is reduced when increasing *n*, due to the increased survival costs that are incurred by expressing increasing numbers of ornaments and preferences. However, average exaggeration of preferences stabilises for increasing *n*, whilst the opposite is true under classical open-ended preferences for high values of *γ*. For the models shown, *a*=1, *λ*=1, *b*=0.0025, *c*=0.5, *μ*_*P*,_ and *μ*_*T*,_ =0.05, *ϑ*=0.2, *u*_*v*_ = 0.9, and each set of parameters was replicated 10 times. Dots represent the endpoints of each individual simulation, and the starting values for ornamentation and preference were *t*=1 and *p*=3. Each data point depicts the grand mean value of *p* (and *t*), which reflects the average value taken over all mean values 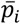 (or 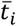) at generation 50,000. Trendlines and 95% confidence intervals are displayed for both positive and negative exaggeration of characters.

## Discussion

Overall, our simulations confirm that including a female Weber-based preference function in models of sexual selection influences the coevolution of male ornaments and female preferences, with the outcome depending on the costs of female choice. Typically, incorporating Weber’s law into female mate choice resulted in less exaggeration of ornaments and preferences compared with open ended preferences. Such reductions in exaggeration of sexually selected characters are particularly likely to happen when female survival does not decrease rapidly if female preferences evolve away from the naturally selected preference optimum (*γ* is low).

In contrast, relative to a standard open-ended preference function, Weber preferences cause a slight exaggeration and diversification of ornaments when survival tapers off rapidly from the naturally selected preference optimum (*γ* is high). Considering males in a population with a female Weber-based preference function, it is relatively straightforward to consider the evolution of male ornamentation away from the naturally selected ornament optimum: males with larger ornaments are selectively favoured in comparison to standard models of sexual selection. Otherwise, the resulting ornament size and variation (noise distribution) perceived by females would overlap with ornament of sizes 0 or less, giving such males little advantage relative to those mated with by females at random. Thus, a considerable ornament investment would be necessary to notably increase a male’s chance of mating.

As with many models of sexual selection, the crux lies in the evolution of female preferences: we would expect that Weber noise would selectively favour and exaggerate larger female preferences relative to standard models of sexual selection, so that females would be better able to distinguish among the perceived noise in male ornaments. However, any such exaggeration of female preferences to larger character values beyond standard models would be immediately checked by any increased costs of female choice. This is indeed the case for scenarios where female survival does not decrease rapidly if female preferences evolve away from the naturally selected preference optimum (*γ* = 1 and *γ* = 2), but for scenarios where survival tapers off rapidly from the naturally selected preference optimum (larger values of *γ*), the fitness landscape is almost flat for female preferences that are only slightly above 0 (see Figure 7). Consequently, smaller preference values can evolve without survival costs, which in turn selectively favours males to evolve ornaments of similar sign away from the naturally selected optimum (*t* = 0). However, in contrast to standard models of sexual selection which would favour only ornaments of modest size, ornaments in Weber contexts are under stronger selection to evolve to larger values to escape the perceived overlap of ornament values with ornament values that are 0 or less. While the resulting exaggeration is only slight (see Figure 5L), it is also not negligible and consistent for larger numbers of ornaments and preferences (e.g., see Figures 3&6). Consequently, for highly nonlinear cost functions, Weber preferences can result in greater exaggeration relative to classical models of sexual selection.

**Figure 7:**
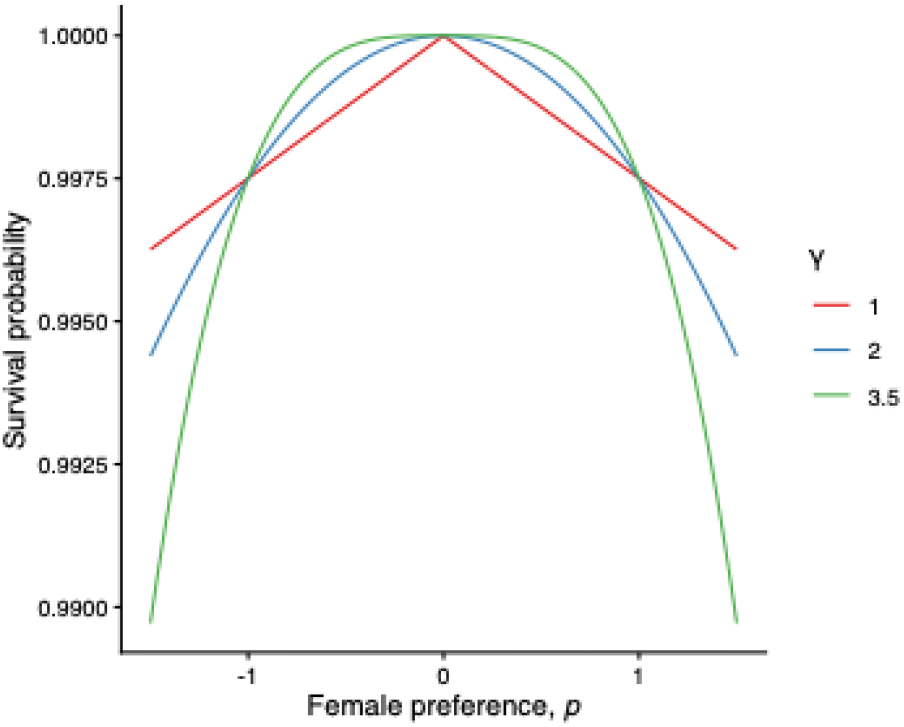
The survival probability away from the naturally selected optimum for different values of *γ*. Parameters: *b* = 0.0025.

To date, there are three main (and until now untested) theoretical arguments on how Weber’s law of proportional processing may shape the coevolution of ornaments and preferences (LaBarbera et al., 2020). Weber processing may 1) slow or halt the evolutionary exaggeration of a male ornament, as the ever-larger increments in ornament value required to elicit a female response become unachievable (Cohen, 1984; Ryan & Cummings, 2013); 2) favour the evolution of more exaggerated male ornaments through chase-away sexual selection, as females require more and more intense stimuli to elicit a response (Holland & Rice, 1998; Ryan & Cummings, 2013), or 3) favour the evolution of novel ornament components to improve signal efficacy once initial ornaments have reached the point of negligible fitness returns on further exaggeration, (Akre & Johnsen, 2016; Endler et al., 2005). In the case of *γ* being low (survival not decreasing rapidly from the naturally selected optimum), scenario (1) occurs, where the decelerating returns of the preference function cause a halt in ornament and preference exaggeration.

However, as survival selection becomes stronger and more of a problem (*γ* is high), the traits in the multidimensional models become more exaggerated and differentiated, and we see more diverse male ornaments evolving when considering Weber’s law. This suggests that Weber preferences favour more diversification of exaggerated characters, at least when the reduction in female survival due to expressing multiple trait preferences is more severe. Indeed, multi-component signals have been shown to outperform single-component signals in discriminability by reducing reaction times, increasing the probability of detection, and lowering the intensity at which detection occurs, especially when components are produced over multiple sensory modalities (Hebets & Papaj, 2005; Rowe, 1999). This could highlight an alternate mechanism to bypass female proportional processing constraints when variation in a single ornament has been exhausted. Analytical models here would be useful to decipher why these trends appear in these individual-based models.

Evidently, if a population or species does adhere to Weber’s law during mate choice, then the coevolution of ornaments and preferences will be impacted, and our understanding of sexual selection and its effects on trait evolution is fundamentally changed. If the cost of a female expressing a preference is low in terms of survival (Martin & Hosken, 2004; Sharma et al., 2010), then we would not expect significant exaggeration in ornaments or preferences, but if the cost is quite dramatic in terms of survival (Engelhard et al., 1989; Friberg & Arnqvist, 2003; Lee et al., 1996; Magnhagen, 1991), then we would expect more exaggerated and diversified male ornaments. This is clearly counterintuitive to the predictions of classical models of sexual selection, and important to highlight. To unravel these consequences effectively, it would be useful to conduct more research on the survival costs of female preferences in the real world. Whilst there has been an abundance of work on the costs of male sexual signalling (Dougherty, 2021; Scharf et al., 2013; White et al., 2022), most theoretical models assume an exact shape of the costs of female preferences, when in reality there is little empirical information about this and how rapidly survival drops off away from a certain survival optimum (Kotiaho, 2001; Kuijper et al., 2012). Ultimately, we need more studies documenting the varying survival costs of expressing a preference.

In conclusion, we demonstrate that including a Weber-based preference function in individual-based models of sexual selection can influence outcomes for the coevolution of male ornaments and female preferences, in both a Fisherian and a good genes scenario. For many years, theoretical work has overlooked Weber’s law of proportional processing when examining sexual selection – indeed, Iwasa and Pomiankowski (1995) even state that ‘small ornaments are worse indicators of male viability because differences in size are more difficult to distinguish’, whilst the opposite is true under Weber processing i.e. it is easier for females to discriminate between smaller ornaments. We show that introducing Weber’s law into models of sexual selection causes the exaggeration of male ornaments and female preferences to stagnate, or exaggerate and diversify across multiple traits, depending on how rapidly survival falls off away from the naturally selected optimum (*γ*). Future theoretical work should therefore aim to consider Weber’s law when studying sexual selection, as it is highly relevant in the real world and can clearly dramatically influence the outcomes of these models. Models investigating sexual conflict, the derivation of a Weber preference function from first principles, and even applying non-linear processing more broadly in a signalling context such as male mate choice and male-male assessment would also be interesting to consider following the results found here.

## Acknowledgements

We would like to thank DH for providing comments that greatly improved the manuscript, and EMC for initial discussion of ideas.

## Data availability statement

Code and instructions on how to run it are publicly available at [https://github.com/kathrynbullough/Weber].

