## Supplementary Materials for "Weber’s law of proportional processing influences coevolution of ornaments and preferences in models of sexual selection"

**S1: Open-ended preferences preclude Weber’s Law**

Focusing on the univariate model of sexual selection, it is very easy to show that open-ended preferences -- for which the odds of mating are given by $\psi\left( T|P \right)=\exp\left( aPT \right)$ -- preclude Weber’s Law and result in a constant mating advantage that is invariant with respect to the ornament size *T*. Consider a female who expresses a preference of strength *P* chooses among two males, in which the first male has an ornament of size $T+\Delta$ while the second male has an ornament of size $T$. The probability $\Pr\left( T+\Delta| P \right)$ that the female chooses the male with ornament size $T+\Delta$ is then given by $\Pr\left( T+\Delta| P \right)=\exp\left[ aP\left( T+\Delta\right) \right]/(\exp\left[ aP\left( T+\Delta\right) \right]+\exp\left( aPT \right))$. Multiplying the rhs by $\exp\left( -aPT \right)/\exp\left( -aPT \right)$ then yields $\Pr\left( T+\Delta| P \right)=\exp\left( ap\Delta\right)/(1+exp \left( ap\Delta\right))$, which is invariant with respect to the magnitude of the ornament *T*. Consequently, in case of open-ended preferences, females are always able to distinguish a difference of $\Delta$ units between the two ornaments, regardless of the overall magnitude of the ornament.

**S2: Occurrence of positive and negative equilibria in Fisherian vs good-genes models**

In the models of good-genes sexual selection considered in this study, exaggerated preferences and ornaments that signal genetic quality appear to settle on two alternative equilibria, in which ornaments and preferences are either positive or negative (Figures 4-6). By contrast, in case of Fisherian sexual selection (Figures 1-3), ornaments and preferences typically settle at positive values. This difference arises from assumptions about the consequences of strictly negative biases on ornaments versus genetic quality (Pomiankowski et al 1991, Iwasa et al 1991). We set out these differences below.

**Strictly negative mutation biases in Fisherian models favour positive equilibria of *t*, *p***

In the Fisherian model by Pomiankowski et al (1991), mutation biases are always directed towards ever more negative values of the ornament (regardless of whether the current value of the ornament is positive or negative) and we have followed the same assumption in the current model. Consequently, due to these negative mutation biases, males with negative ornament values are more likely to arise than males with positive ornaments, at least when the population is close to the naturally selected optimum of *t* = 0, and more so when *t* is negative. Consequently, the prevalence of males with negative ornaments reduces the indirect benefit of a female who expresses preferences for males with negative ornaments, as her sons are more likely to face competition from males with similar or even more negative values of *t*. By contrast, sons born from a female with a preference for positive ornaments are less likely to suffer increased competition from other males with positive ornaments. This is because such males are less likely emerge, because of a negative mutation bias reducing values of ornaments. Consequently, because indirect benefits are larger when expressing preferences for positive-valued ornaments, female preferences are more likely to favour positive ornaments. Hence, coevolution is more likely to result in a positive equilibrium of *t* and *p*.

To assess this effect of a negative mutation bias acting on the ornament, we have also considered another type of mutation bias in which mutations are opposite in sign to the current value of the ornament (Figure S2.1). Such a mutation bias reduces the magnitude of the ornament towards 0, but removes the artefact that males with negative ornaments are more likely to emerge than males with positive ornaments when *t* ≤ 0. In this case exaggerated ornaments and preferences are now indeed equally likely to occur at positive or negative equilibria, unless mutation biases are very large.

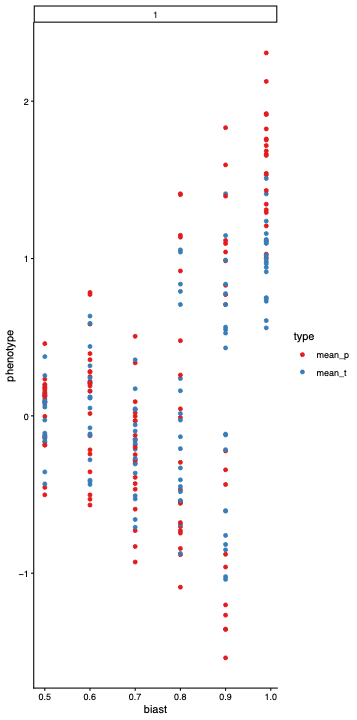

**Figure S2.1.** When considering mutation biases that reduce the absolute magnitude rather than mutation biases that are strictly negative, we find that positive and negative equilibria are equally likely in Fisherian models of sexual selection.

**Strictly negative mutation biases in good-genes models result in positive or negative *t*, *p* equilibria**

The action of the mutation bias is different in good-genes models: first, the strictly negative mutation bias *u_v_* now acts on the genetic quality locus *v*, whereas mutations on *t* are not biased. Moreover, ornaments are now given by the product *tv*, reflecting a condition-dependent trait. Because the optimal genetic quality *v*_opt_ is set at a large and positive value in both classical models and the current study (*v*_opt_ = 10 in Iwasa et al 1991 and in the current study), negative values of *v* are highly unlikely to occur, as values *v* < *v*_opt_ are rapidly removed by natural selection. Hence, because typically *v* > 0, should a male with a negative ornament value *tv* < 0 arise, this is not a result of strictly negative biased mutations on *v*, but rather due to unbiased mutations acting on *t* that result in *t* < 0. Consequently, in good-genes models, because *v* is large and positive, mutation biases are not making it more likely that negative values of ornamentation occur, in contrast to the Fisherian model above. Instead, in good-genes models, males with negative ornaments are equally likely to arise as males with positive ornaments are. Hence, a female in a good-genes model stands to gain similar indirect benefits by having preferences for negatively ornamented males versus preferences for positively ornamented males. This then explains why exaggeration in a good-genes model can result either in an equilibrium of *t*, *p* > 0 or an equilibrium of *t*, *p* < 0.

**S3: Supplementary figures**

**Fisherian sexual selection**

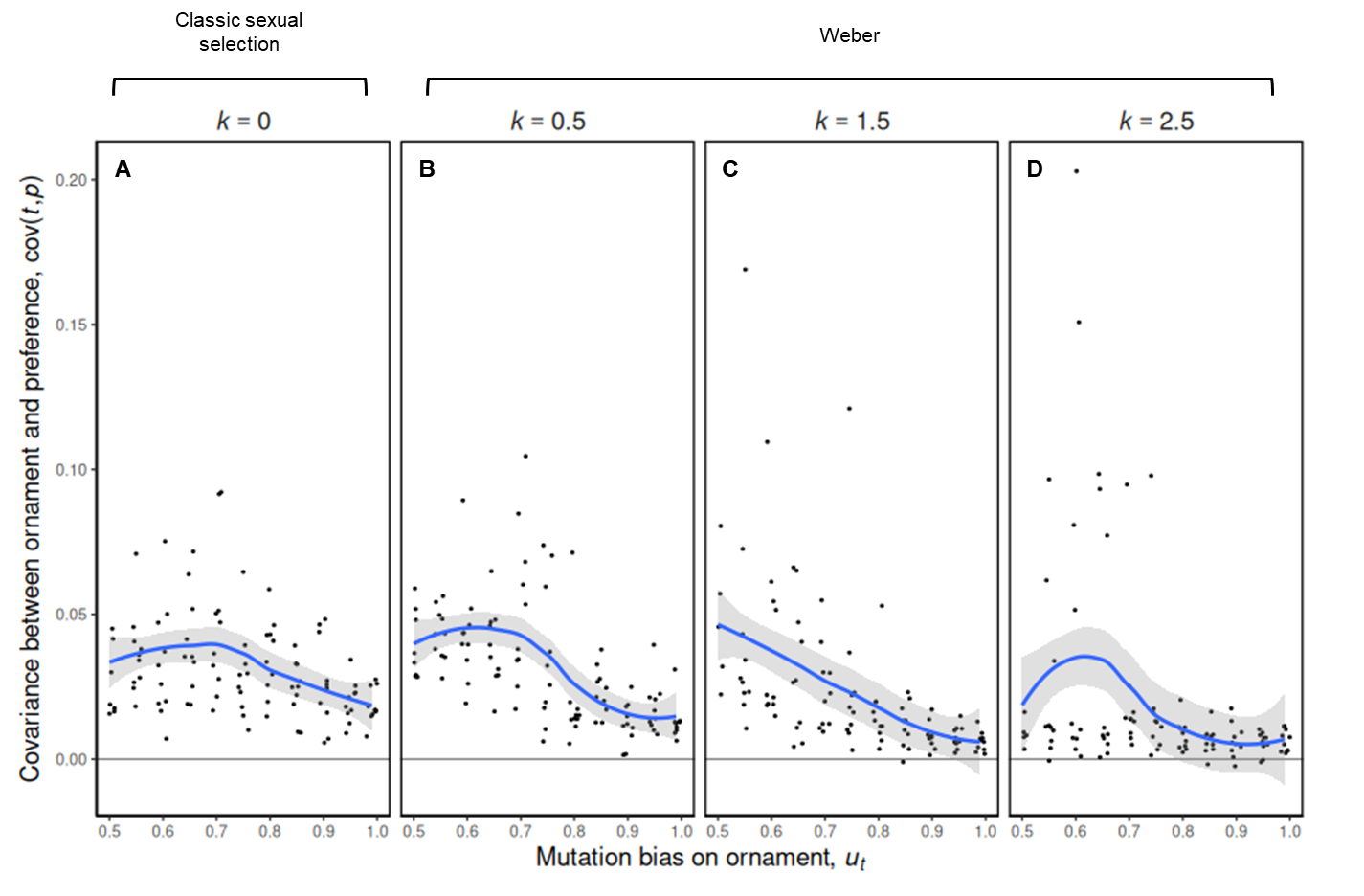

**Figure S3.1**: A unidimensional individual-based model of Fisherian sexual selection, showing the covariance between one male ornament (*t*) and one female preference (*p*) as they coevolve. For all scenarios, as deleterious mutation bias on the male ornament increases, covariance between ornaments and preferences decreases, and this is more pronounced under Weber preferences. Parameters: *a*=1, *λ*=1, *b*=0.0025, *c*=0.5, *μ_P,_* and *μ_T,_* =0.05, *γ*=2, and 10 replicate simulations for each unique parameter combination. The starting values for ornamentation and preference were *t*=1 and *p*=3.

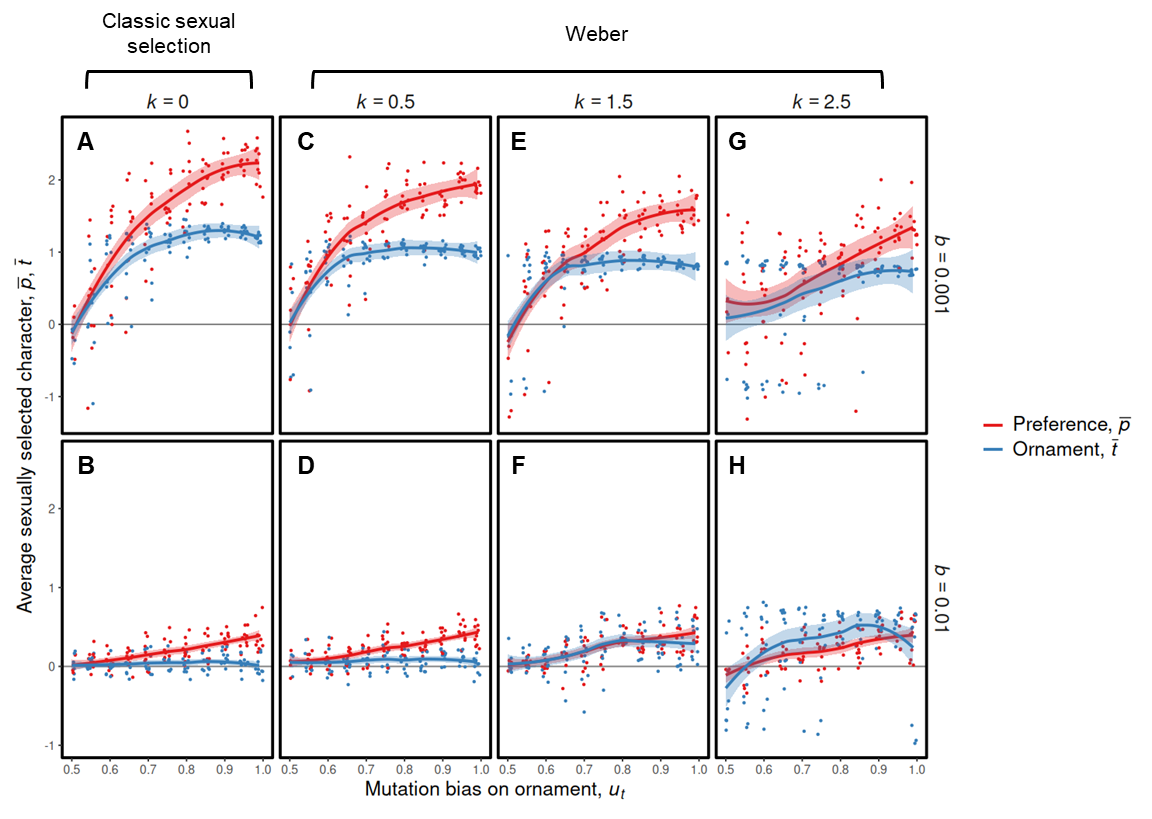

**Figure S3.2**: A unidimensional individual-based model of Fisherian sexual selection, varying the strength of Weber preferences *k* and the cost of choice *b*. As the deleterious mutation bias *u_t_* on the male ornament increases, exaggeration of ornaments and preferences also increases, yet with increasing *k* and *b* exaggeration of ornaments and preferences moderately reduces. Parameters: *a*=1, *λ*=1, *c*=0.5, *μ_P,_* and *μ_T,_* =0.05, *γ*=2, and 10 replicate simulations for each unique parameter combination. The starting values for ornamentation and preference were *t*=1 and *p*=3.

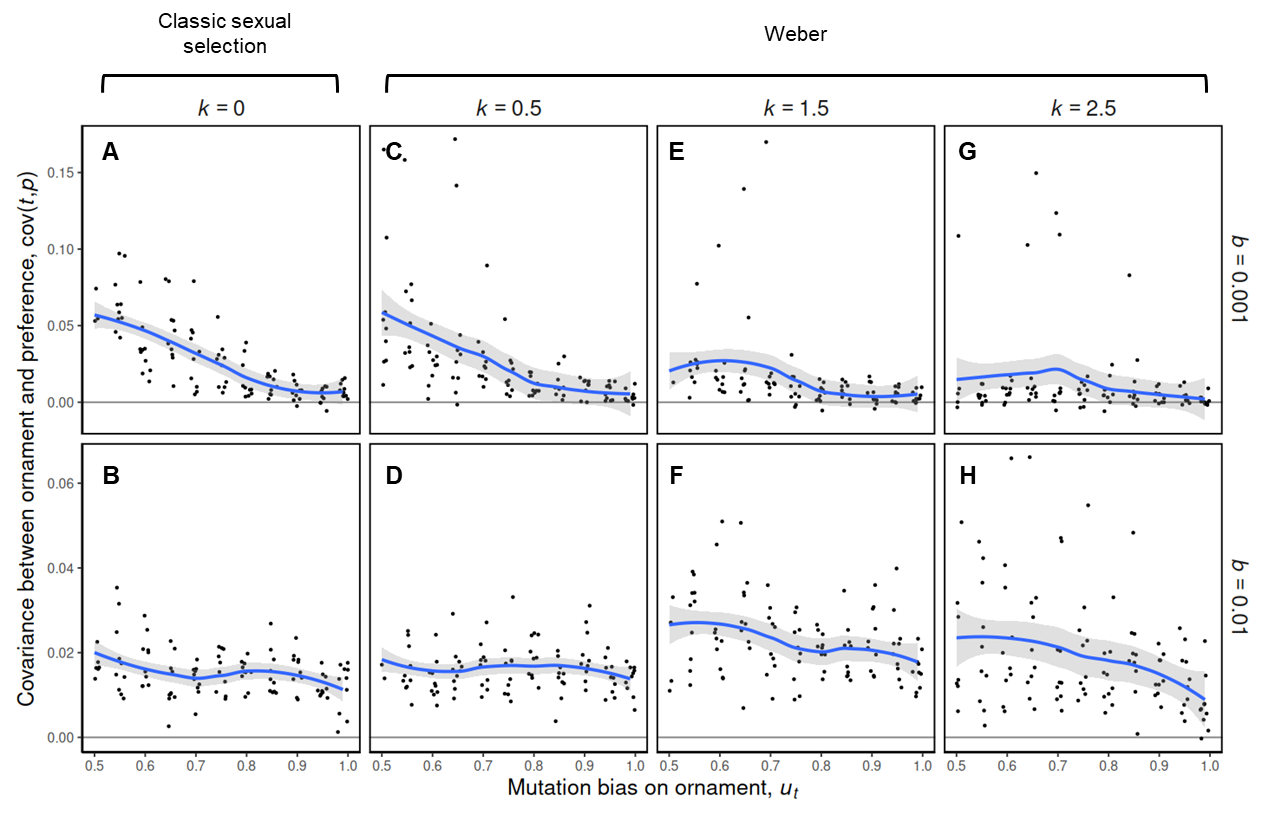

**Figure S3.3**: A unidimensional individual-based model of Fisherian sexual selection, showing the covariance between one male ornament (*t*) and one female preference (*p*) as they coevolve whist varying the strength of Weber preferences *k* and the cost of choice *b*. For all scenarios, as deleterious mutation bias on the male ornament increases, covariance between ornaments and preferences decreases, and this is moderately more pronounced under Weber preferences and smaller values of *b*. Parameters: *a*=1, *λ*=1, *c*=0.5, *μ_P,_* and *μ_T,_* =0.05, *γ*=2, and 10 replicate simulations for each unique parameter combination. The starting values for ornamentation and preference were *t*=1 and *p*=3.

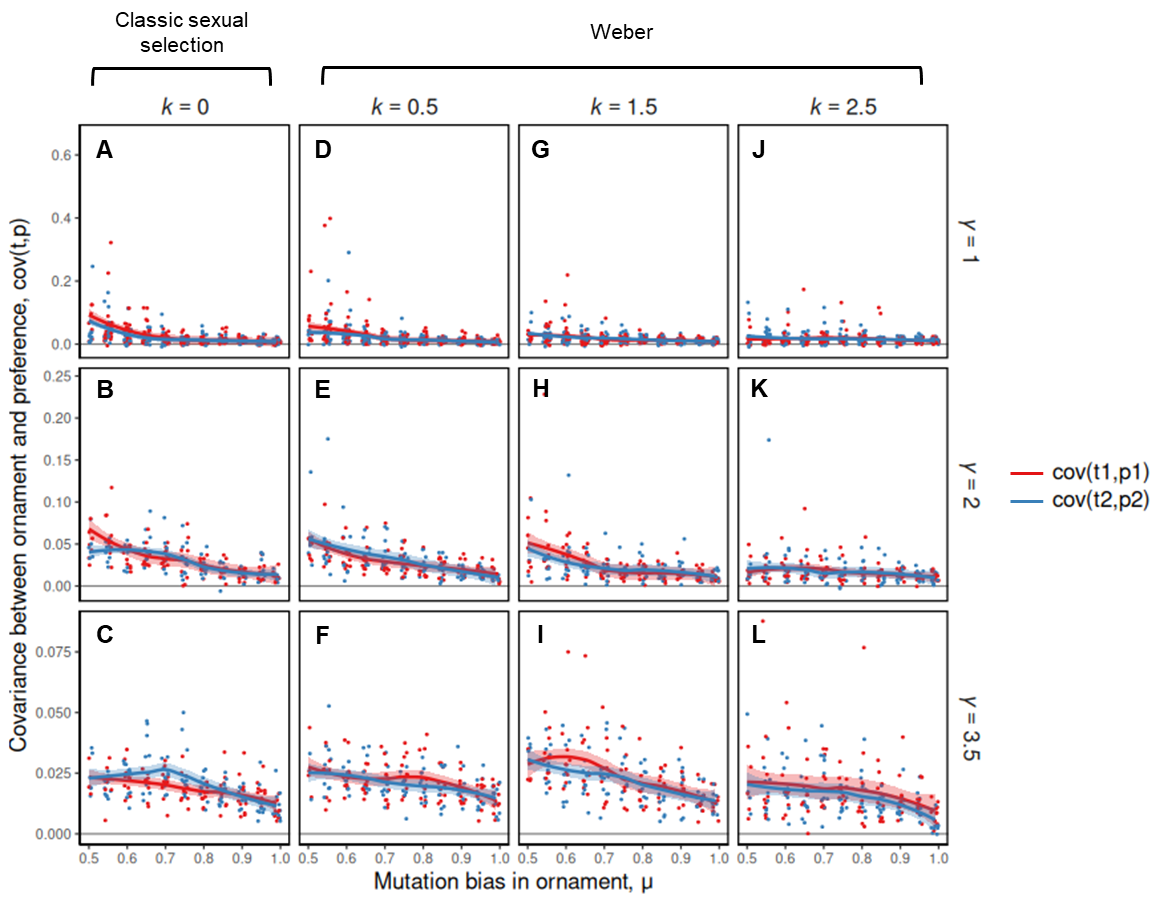

**Figure S3.4**: A multidimensional individual-based model of Fisherian sexual selection, showing the covariance between two male ornaments (*t1* and *t2*) and two female preferences (*p1* and *p2*) as they coevolve. For all scenarios, as deleterious mutation bias on the male ornaments increase, covariance between ornaments and preferences decreases, and this is more pronounced under the Weber preferences with low values of gamma. Under the Weber preferences with high values of gamma, this is less pronounced. For the models shown, *a*=1, *λ*=1, *b*=0.0025, *c*=0.5, *μ_P_*_,_ and *μ_T_*_,_ =0.05, *ϑ*=0.2, and each set of parameters was replicated 10 times. The starting values for ornamentation and preference were *t*=1 and *p*=3.

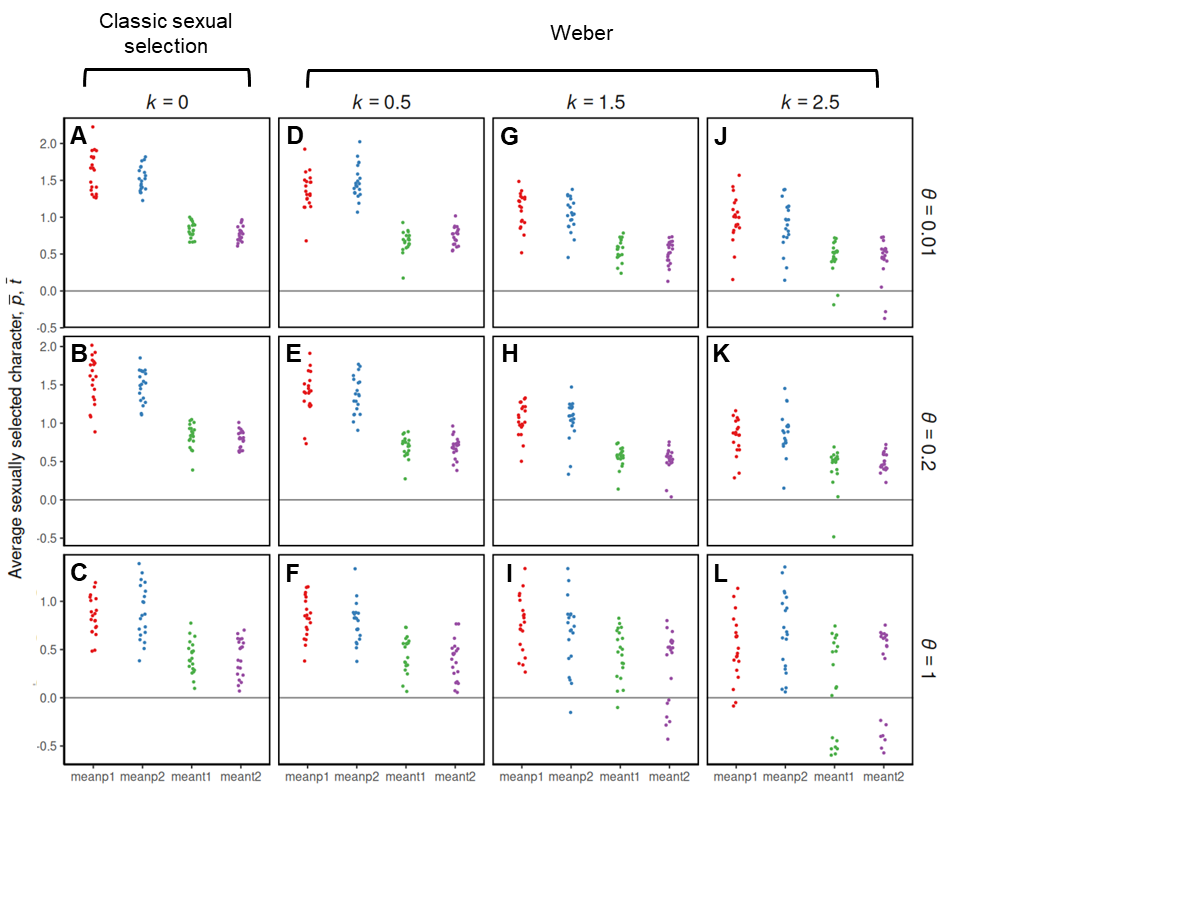

**Figure S3.5**: A multidimensional individual-based model of Fisherian sexual selection, showing equilibrium exaggeration of two male ornaments (*t1* and *t2*) and two female preferences (*p1* and *p2*) at generation 150,000. Exaggeration is less pronounced under Weber preferences and for high values of *ϑ*. For the models shown, *a*=1, *λ*=1, *b*=0.0025, *c*=0.5, *μ_P_*_,_ and *μ_T_*_,_ =0.05, *γ*=2, *u_t_* = 0.99, and each set of parameters was replicated 20 times. The starting values for ornamentation and preference were *t_1_*=*t_2_*=1 and *p_1_*=*p_2_*=3.

**Good genes sexual selection**

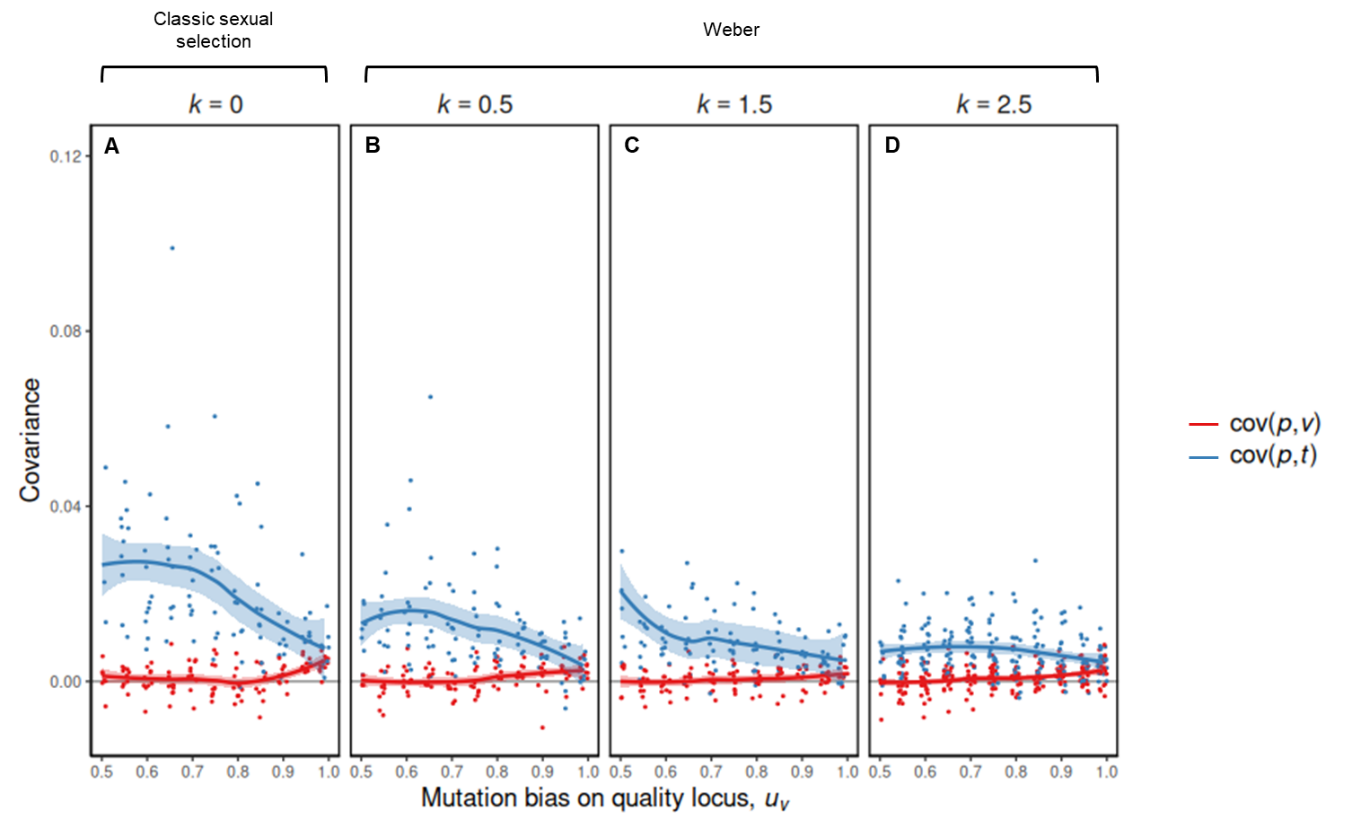

**Figure S3.6**: A unidimensional individual-based model of good genes sexual selection, showing the covariance between one female preference (*p*), and either one male ornament (*t*) or one male quality locus (*v*) as they coevolve. For all scenarios, as deleterious mutation bias on the male quality locus increases, covariance between ornaments and preferences decreases, and this is more pronounced under Weber preferences. Covariance between *p* and *v* remains low under all scenarios. For the models shown, *a*=1, *λ*=1, *b*=0.0025, *c*=0.5, *μ_P_*_,_ and *μ_T_*_,_ =0.05, *γ*=2, and each set of parameters was replicated 10 times. The starting values for ornamentation and preference were *t*=1 and *p*=3.

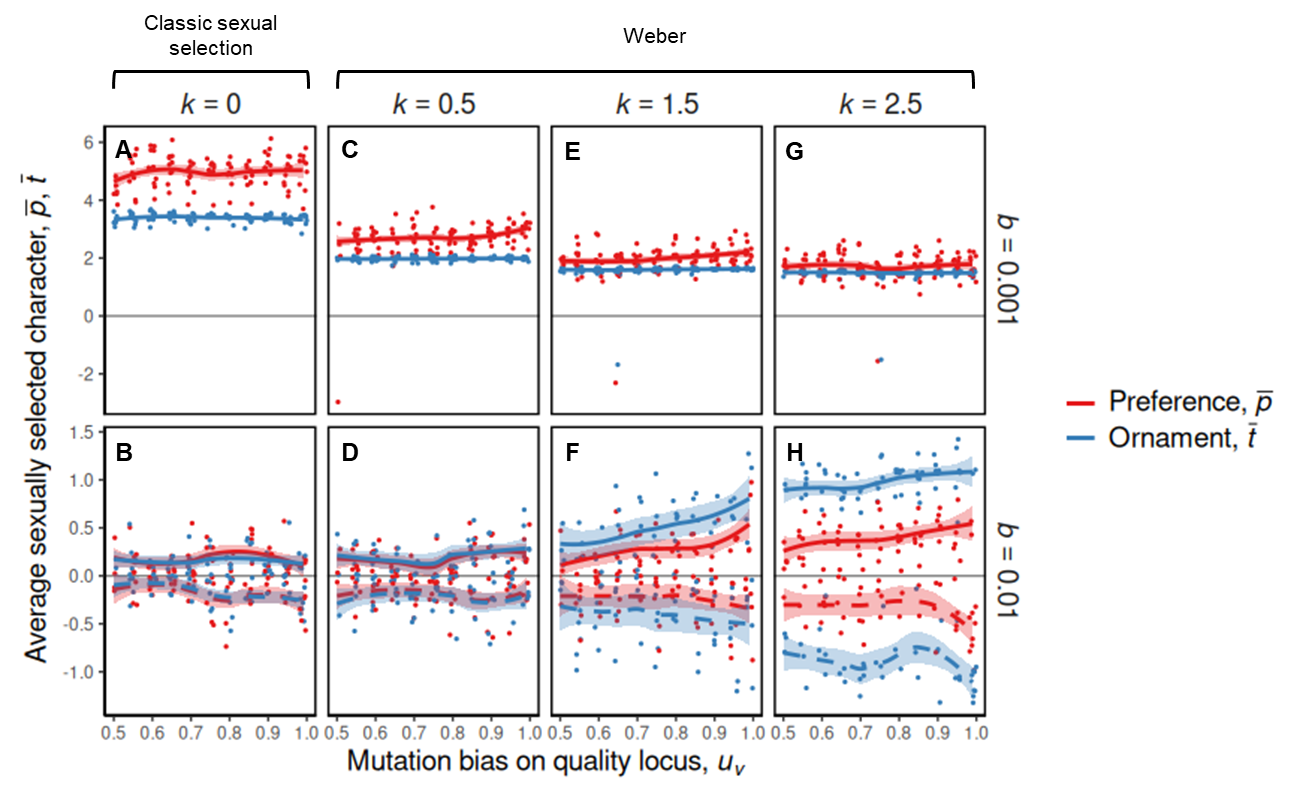

**Figure S3.7**: A unidimensional individual-based model of good genes sexual selection, varying the strength of Weber preferences *k* and the cost of choice *b*. For high values of *b*, exaggeration of ornaments and preferences is notably reduced, however to a lesser extent under stronger Weber preferences. For the models shown, *a*=1, *λ*=1, *c*=0.5, *μ_P_*_,_ and *μ_T_*_,_ =0.05, *γ*=2, and each set of parameters was replicated 10 times. The starting values for ornamentation and preference were *t*=1 and *p*=3.

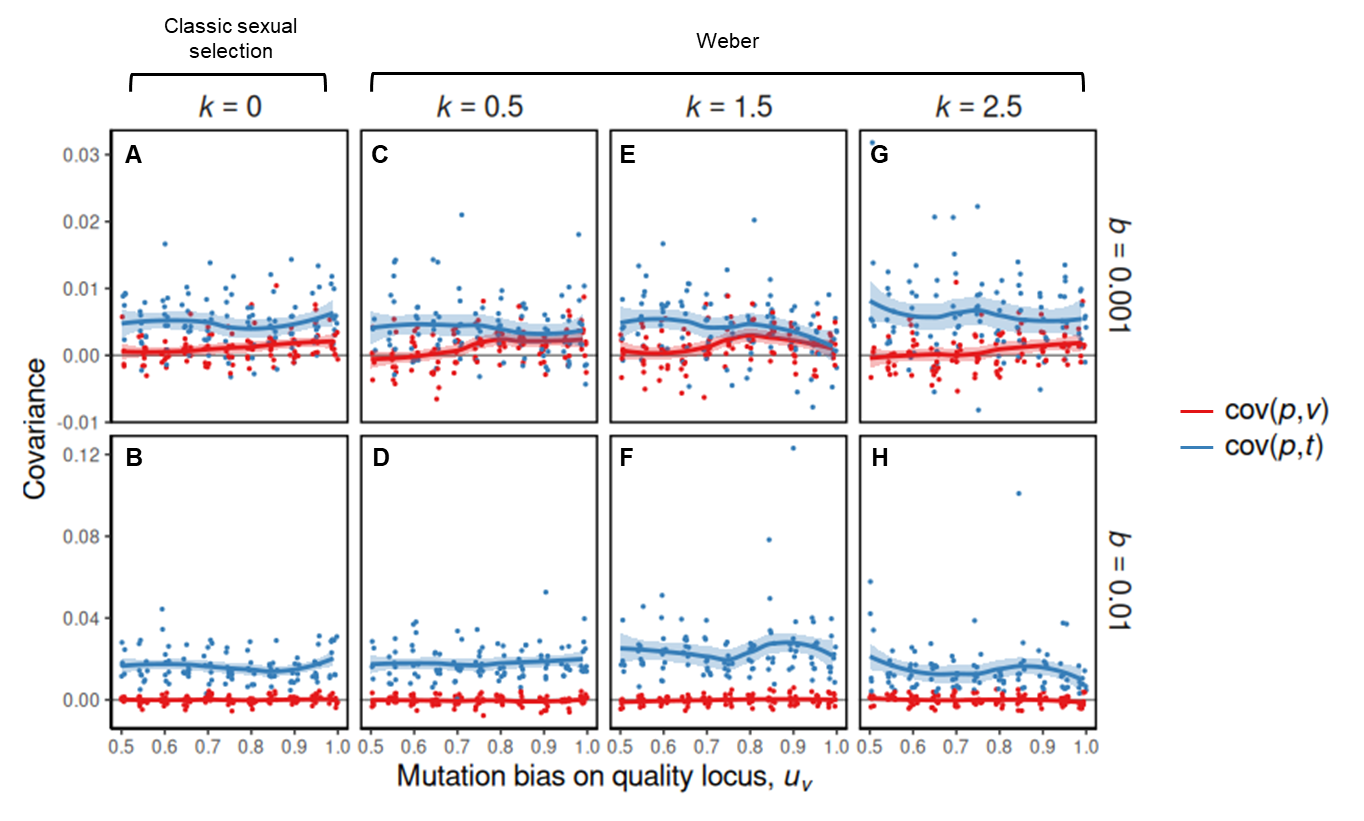

**Figure S3.8**: A unidimensional individual-based model of good genes sexual selection, showing the covariance between one female preference (*p*), and either one male ornament (*t*) or one male quality locus (*v*) as they coevolve, whist varying the strength of Weber preferences *k* and the cost of choice *b*. For all scenarios, covariance between ornaments and preferences is reduced under lower values of *b*, with these trends remaining under varying strengths of Weber preferences. Covariance between *p* and *v* remains low under all scenarios. Parameters: *a*=1, *λ*=1, *c*=0.5, *μ_P,_* and *μ_T,_* =0.05, *γ*=2, and 10 replicate simulations for each unique parameter combination. The starting values for ornamentation and preference were *t*=1 and *p*=3.

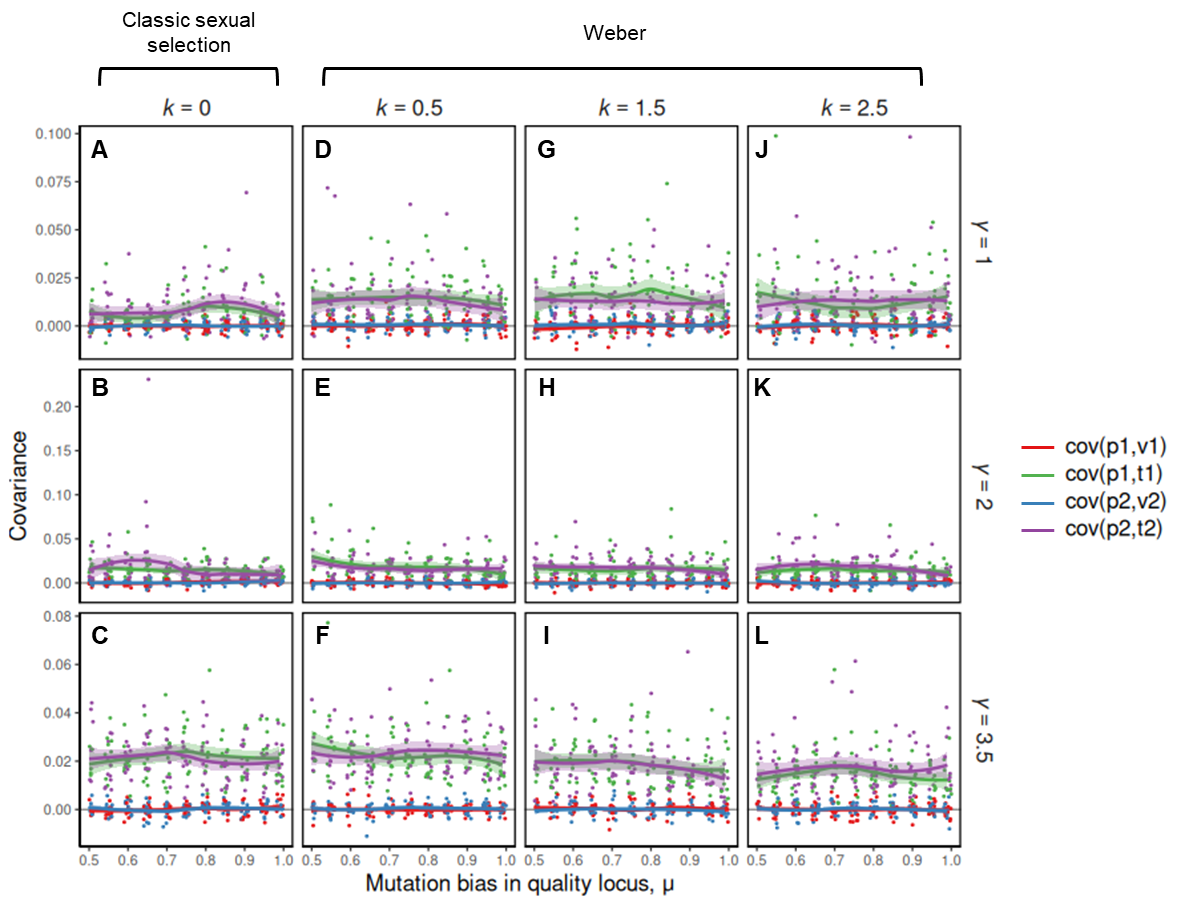

**Figure S3.9**: A multidimensional individual-based model of good genes sexual selection, showing the covariance between two female preferences (*p1* and *p2*), and either two male ornaments (*t1* and *t2*) or two male quality loci (*v1* and *v2*) as they coevolve. Covariance remains low and constant under all scenarios. For the models shown, *a*=1, *λ*=1, *b*=0.0025, *c*=0.5, *μ_P_*_,_ and *μ_T,_* =0.05, *ϑ*=0.2, and each set of parameters was replicated 10 times. The starting values for ornamentation and preference were *t_1_*=*t_2_*=1 and *p_1_*=*p_2_*=3.

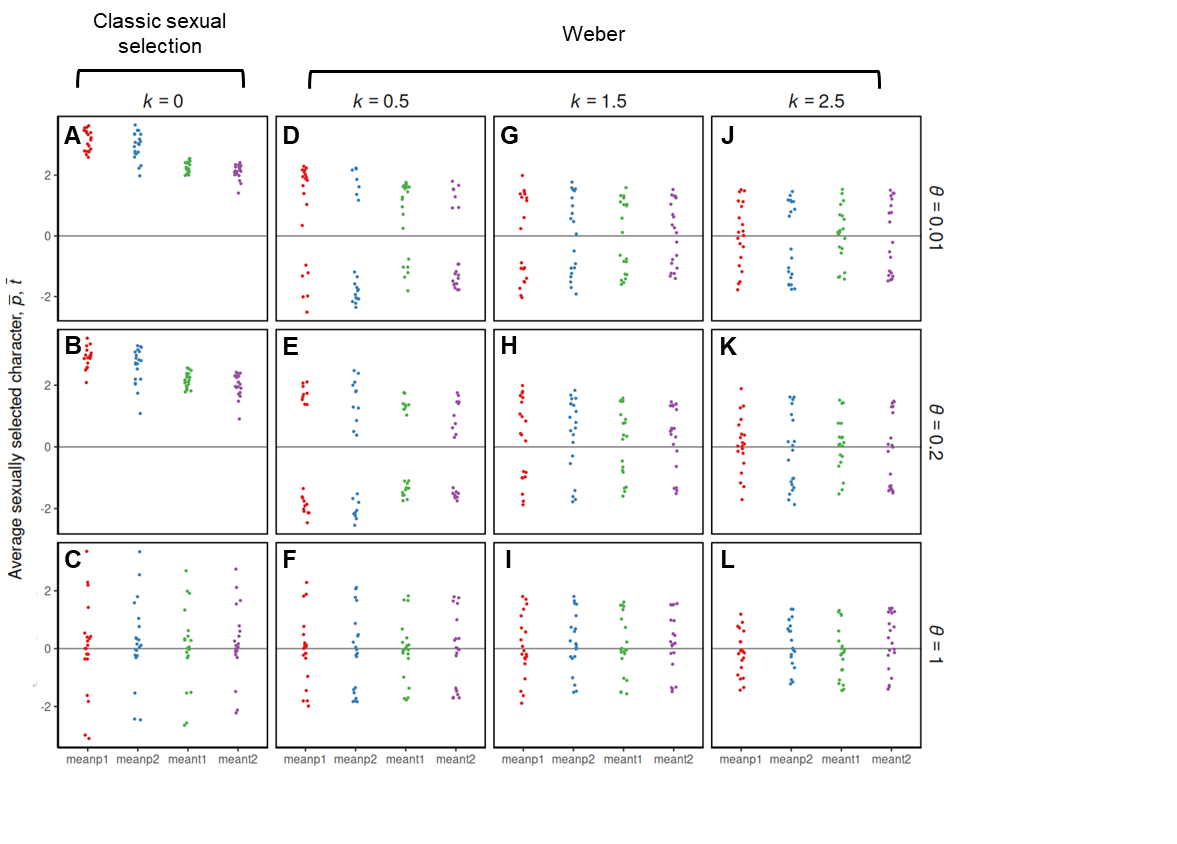

**Figure S3.10**: multidimensional individual-based model of good genes sexual selection, showing coevolution of two male ornaments (*t_1_* and *t_2_*) and two female preferences (*p_1_* and *p_2_*) at generation 50,000 for a scenario in which biased mutations reducing genetic quality occur with probability *u_v_* = 0.99. Exaggeration is less pronounced under Weber preferences and for high values of *ϑ*. For the models shown, *a*=1, *λ*=1, *b*=0.0025, *c*=0.5, *μ_P_*_,_ and *μ_T,_* =0.05, *γ* =2, and each set of parameters was replicated 20 times. The starting values for ornamentation and preference were *t_1_*=*t_2_*=1 and *p_1_*=*p_2_*=3.
